# Whitefly effector G4 interacts with tomato proteins of which MIPDB141 affects whitefly performance

**DOI:** 10.1101/2023.03.11.532171

**Authors:** Diana Naalden, Wannes Dermauw, Aris Ilias, Geert Baggerman, Marieke Mastop, Sarmina Dangol, Nicolas Gaertner, Winfried Roseboom, Mark Kwaaitaal, Gertjan Kramer, Harrold A. van den Burg, John Vontas, Thomas Van Leeuwen, Merijn R. Kant, Robert C. Schuurink

## Abstract

The phloem-feeding insect *Bemisia tabaci* is an important pest, responsible for the transmission of several crop-threatening virus species. While feeding, the insect secretes a cocktail of effectors to modulate defense responses. Here, we present a set of proteins that was identified in artificial diet on which *B. tabaci* was salivating. We studied whether these candidate effectors can play a role in plant immune suppression. Effector G4 was the most robust suppressor of the flg22-induced ROS response when transiently expressed in *Nicotiana benthamiana*. In addition, G4 was able to supress ROS in *Solanum lycopersicum* (tomato) and *Capsicum annuum* (pepper). Fused to a fluorescence tag, G4 localized in the cytoplasm in *N. benthamiana*. A yeast two-hybrid screen combined with a luciferase bimolecular complementation and co-localization assays resulted in the identification of two target proteins in tomato: REF-like stress related protein 1 (RSP1) and meloidogyne-induced giant cell protein DB141 (MIPDB141). Silencing of *MIPDB141* in tomato, using virus-induced gene silencing, reduced whitefly fecundity up to 40% demonstrating that the protein is involved in susceptibility to *B. tabaci*. Together our data demonstrate that effector G4 impairs tomato immunity to whiteflies by interfering with the ROS production and via a direct interaction with tomato susceptibility protein MIPDB141.

## Introduction

The silverleaf whitefly, *Bemisia tabaci*, is a priority pest globally. It has a broad host range and the damage it inflicts directly and indirectly cause tremendous economic losses in agriculture (Saurabh et al., 2021). Direct damage is caused by the insect’s phloem feeding while the major indirect damage is a result of the transmission of many plant damaging viruses (reviewed in Fiallo-Olivé et al. (2020)) but also since the honeydew it secretes attracts secondary plant damaging organisms. *Bemisia tabaci* comprises a species complex of more than 40 cryptic species, with variation in for example host range, insecticide resistance, virus transmission, morphometrics and genetics (De Barro et al., 2011; Mugerwa et al., 2018; Mugerwa et al., 2021). Within this species complex, Mediterranean (MED, also known as the Q biotype) and Middle-East-Asia Minor 1 (MEAM1, also known as the B biotype or *Bemisia argentifolii*) are among the best studied (Boykin et al., 2013).

Whiteflies are herbivores and are in close contact with their host and depend on them throughout their lifecycle. The feeding stages of the insect penetrate the leaf tissue via the apoplast using their stylets to feed from the contents of the sieve tubes of the phloem. Eggs are deposited on the abaxial side of the leaf and after 5-9 days these hatches, after which the mobile juveniles (crawlers) start searching for a suitable feeding site. After selecting a feeding site, crawlers then molt into nymphs that are sessile. The immobile nymphs feed on a single feeding site where they develop over several weeks (depending on environmental conditions) through four nymphal stages into adults (Gangwar and Gangwar, 2018). During this interaction the plant can execute a set of physio-metabolic programs aimed at resisting whitefly feeding, for example via the production of acylsugars in type IV glandular trichomes (Rodríguez-López et al., 2012) and by increasing cuticle thickness (Firdaus et al., 2011). These resistance factors are effective to repel whiteflies yet often insufficient to reduce damage to levels acceptable for crops (Naalden et al., 2021). Pesticides are often used to control whiteflies but this can have undesirable impacts on environment and human health. In addition, frequent use of pesticides against *B. tabaci* often selects quickly for resistance (Biswajit Patra, 2022; Horowitz et al., 2020). Biological control, for example by means of entomopathogenic fungi or parasitoids, can be effective yet its applicability is limited across different environmental conditions and obtaining control takes a relatively long time (Liu et al., 2015; Sani et al., 2020). Therefore, generation of varieties resistant to whiteflies is an important strategy for disease management. To date, only one resistance gene, *Mi-1*.*2* provides enhanced resistance against both MED and MEAM1 biotypes in tomato. However, *Mi-1*.*2* does not provide resistance at higher temperature and in younger tomato plants (Nombela et al., 2003) and thus its applicability is limited.

Whitefly feeding is rapidly detected by plants triggering the plant’s immune response. Yet these responses often are insufficient to protect the plant. Research has shown that this inefficiency of the naturally induced responses often is due to unknown salivary substances (often proteins) secreted during feeding and dampening the induced defence response to levels the whitefly can cope with. Such salivary effectors thus can benefit the insect during feeding by restraining the plant’s natural resistance thus preserving the plant’s value as food (Naalden et al., 2021). Some of these effectors have been identified and characterized for *B. tabaci* (Du et al., 2022; Lee et al., 2018; Peng et al., 2023; Su et al., 2019; Wang et al., 2019; Xu et al., 2019). However, for successfully utilizing these effectors to identify and modify their targets for resistance breeding (Tyagi et al., 2020) but also to limit the chances for whitefly adaptations to such novel resistances to emerge, it is important get a comprehensive overview of effector diversity and, subsequently, target diversity. In this study, we identified a set of candidate effectors that were present in the saliva of *B. tabaci* (MED) and characterize one of these, named G4, further for its *in planta* targets and its effect on plant immunity.

## Materials and Methods

### Rearing *Bemisia tabaci*

The *B. tabaci* strain (MED, Q-biotype) used for saliva collection, was initially sampled from greenhouse-cultivated eggplants in May 2019 in Ierapetra, South Crete (Greece) and subsequently maintained on cotton plants in a plant growth cabinet at 25 (±1)°C, 50-60% relative humidity and a 16h/8h light-dark regime at the insectary of the Biology Department, University of Crete (Greece). For isolation of effectors and bioassays the *B. tabaci* population (MEAM1) was maintained on cucumber plants in insect cages in the greenhouse at 28°C under 16h/8h light-dark regime at the University of Amsterdam.

### *Bemisia tabaci* saliva collection

Petri dishes (55 mm diameter) with two ventilation holes covered with thin mesh on the side walls, were used as feeding chamber. Approximately, two hundred newly emerged whitefly females were immobilized using carbon dioxide and placed in a Petri dish. The Petri dish was covered by two layers of parafilm with 1 mL of artificial diet containing 30% sucrose and rifampicin (0.05 mg/ml) between the two layers. The feeding chambers were placed at 25 (±1)°C, 50-60% relative humidity and a 16h/8h light-dark regime in a growth cabinet. Three biological replicates of artificial diet enriched with whitefly saliva were collected (approximately 800 μl of artificial diet/replicate) using a micropipette after six and twenty-four hours of whitefly feeding (Btab_6h or Btab_24h). Three replicates of artificial diet without whitefly saliva were also collected at the same time points and served as controls (Con_6h or Con_24h). The entire process was carried out as aseptically as possible. Collected artificial diet samples were kept at -80°C until further use.

### LC-MS/MS analyses of artificial diet

Artificial diet samples were lysed using 200 μl RIPA lysis buffer (ThermoFisher Scientific, Belgium) and 1x HALT protease inhibitor (Thermo Fisher Scientific), combined with a 30 sec during sonication (Branson Sonifier SLPe ultrasonic homogenizer, Labequip, Ontario, Canada) with an amplitude of 50% on ice. After centrifugation of the samples for 15 min at 10,000 g at 4°C, the pellet was discarded. One μg equivalent of proteins was then reduced using 2 μl of 200 mM tris (2-carboxyethyl) phosphine, in a volume of 20 μl 200 mM triethylammonium bicarbonate (TEAB), and incubated for 1 h at 55°C. After alkylation of the proteins with 2 μL of 375 mM iodoacetamide for 30 min protected from light, 6 volumes of ice-cold acetone were added, and the samples were incubated overnight at -20°C. The next day, samples were centrifuged for 10 min at 10.000 g at 4°C, the acetone layer was removed with a pipette, samples were dried using a vacuum centrifuge, and pellets were resolved in 20 μl of 200 mM TEAB. Proteins were then digested with trypsin (Promega) overnight at 37°C with an enzyme trypsin ratio of 1:50. Before LC-MS/MS analysis, the samples were desalted with Pierce C18 spin columns according to the manufacturer’s instructions (Thermo Fisher Scientific).

Each of the digested samples was separated by nano reverse phase C18 (RP-C18) chromatography on a Waters nano-acquity nano-UPLC system using an ACQUITY UPLC M-Class Symmetry C18 Trap Column, 5 μm, 180 μm × 20 mm connected to a ACQUITY UPLC M-Class Peptide BEH C18, 300Å, 1.7 μm, 100 μm × 100 mm analytical column (Waters, USA). Of each sample, a total of 1 μg of peptides were loaded on the column. Before loading, pellet of the digested proteins was dissolved in mobile phase A, containing 2% acetonitrile. A linear gradient of mobile phase B (0.1% formic acid in 100% acetonitrile) from 0 to 45% in 90 min, followed by a steep increase to 100% mobile phase B in 10 min, was used at a flow rate of 300 nL/min. Liquid Chromatography (LC) was connected to a Q-Exactive Plus mass spectrometer equipped with a nanospray ion source (Thermo Fisher, Waltham, MA, USA). The high-resolution mass spectrometer was set up in an MS/MS mode where a full scan spectrum (350 – 1850 m/z, resolution 70,000) was followed by a high energy collision activated dissociation (HCD) tandem mass spectra (100 – 2000 m/z, resolution 17,500). Peptide ions were selected for further interrogation by tandem MS as the twenty most intense peaks of a full scan mass spectrum. The normalized collision energy used was set at 28%. A dynamic exclusion list of 20 sec for the data-dependent acquisition was applied.

### Proteomic data analysis

Peak lists obtained from MS/MS spectra were identified using OMSSA version 2.1.9 (Geer et al., 2004) X!Tandem version X! Tandem Vengeance (2015.12.15.2) (Craig and Beavis, 2004) and MS-GF+ version Release (v2018.04.09) (Kim and Pevzner, 2014). The search was conducted using SearchGUI (version 3.3.16) (Vaudel et al., 2011). Protein identification was conducted against a concatenated target/decoy (Elias and Gygi, 2010) version of the *B. tabaci* MED protein database holding 20,748 target sequences (version of September 2019, downloaded from http://www.whiteflygenomics.org, Data S1, see Supporting Information) and the common Repository of Adventitious Proteins (cRAP) database (116 sequences, available at http://www.thegpm.org/crap/). The decoy sequences were created by reversing the target sequences in SearchGUI. The identification settings were as follows: Trypsin, Specific, with a maximum of 2 missed cleavages; 10.0 ppm as MS1 and 0.5 Da as MS2 tolerances; variable modifications: Carbamidomethylation of C (+57.021464 Da), Oxidation of M (+15.994915 Da), Pyrolidone from E (--18.010565 Da), Pyrolidone from Q (--17.026549 Da), Pyrolidone from carbamidomethylated C (--17.026549 Da), Acetylation of protein N-term (+42.010565 Da), variable modifications during refinement procedure: Pyrolidone from E (--18.010565 Da), Pyrolidone from Q (--17.026549 Da), Pyrolidone from carbamidomethylated C (--17.026549 Da). Peptides and proteins were inferred from the spectrum identification results using PeptideShaker version 1.16.42 (Vaudel et al., 2015). Peptide Spectrum Matches (PSMs), peptides and proteins were validated at a 1.0% False Discovery Rate (FDR) estimated using the decoy hit distribution. The Peptideshaker identification results for each replicate were exported as an excel file (“Default protein report”, Table S1, see Supporting Information) and summarized into one table for each timepoint (6h or 24h of feeding, Table S2, see Supporting Information). Only those proteins that were 1) not cRAP proteins, 2) identified in at least two out of three biological replicates of each treatment (Btab_6h/Btab_24h) and 2) were not identified in any control sample (Con_6h/Con_24h), were regarded as present and reliably identified. The mass spectrometry data along with the identification results have been deposited to the ProteomeXchange Consortium (Vizcaíno et al., 2014) via the PRIDE partner repository (Martens et al., 2005) with the dataset identifiers PXD033180 and 10.6019/PXD033180. Last, the presence of a signal peptide in the detected *B. tabaci* proteins was predicted using SignalP 6.0 (Teufel et al., 2022).

### Sequence analysis of G4

The coding region of G4 was used in a BLAST search against the Whitefly Genome Database (MEAM1; http://www.whiteflygenomics.org/cgi-bin/bta/index.cgi; (Chen et al., 2016)) and the non-redundant protein and nucleotide databases for all organisms (Highly similar sequences (megablast)) of the National Center for Biotechnology Information (NCBI). Alignments of proteins were made using the ClustalW multiple alignment algorithm in Bioedit sequence alignment editor (Hall, 1999). Prediction of cellular localization was performed using Wolf PSort (https://wolfpsort.hgc.jp) and LOCALIZER (https://localizer.csiro.au; Sperschneider et al., 2017) using the mature G4 sequence (without signal peptide) as input. Prediction of domains was performed using InterPro (https://www.ebi.ac.uk/interpro/; Paysan-Lafosse et al., 2023).

### Plant growth

*Nicotiana benthamiana* seeds were germinated in universal soil at 21°C 16h/8h light-dark regime and circa 1-week-old individual plantlets were transferred to pots with universal soil. The plantlets were grown for 4-5 weeks prior to Agrobacterium infiltration. Seeds of *Solanum lycopersicum* (cv. Moneymaker) and *Capsicum annuum* (cv. Mandy) were germinated in universal soil and circa 1-week-old individual plantlets were transferred separately to pots with universal soil. The plantlets were grown for 4-5 weeks at 21°C under light/dark cycles of 16h/8h at 65% relative humidity until used for Reactive Oxygen Species (ROS) assays, phloem extraction and bio-assays. For the VIGS assays, 9 days old *S. lycopersicum* plantlets were agroinfiltrated and kept under the same conditions post agroinfiltration.

### Effector and plant protein constructs

Full-length coding sequences of the candidate effectors were PCR amplified from *B. tabaci* MEAM1 cDNA (adult stage; G1-transcriptome (G1-t), G3 and G4) using gene-specific primers (primers are listed in Table S4, see Supporting information). PCR was performed using VWR® Taq DNA-polymerase with 28 cycles at an annealing temperature of 58 °C and an elongation time of 90 sec. Obtained PCR products were purified according to the manufacturer’s instructions (Genejet PCR Purification Kit, ThermoFisher Scientific) and ligated into the pJET vector according to the manufacturer’s protocol (ThermoFisher Scientific). The plasmids were transformed in *Escherichia coli* (DH5-α) using heat shock treatment (42 sec at 42°C, incubation for 2 min on ice and recovery for 1,5 h in 500 μL LB medium with shaking at 200 rpm at 37°C). Transformed cells were grown overnight (o/n) at 37°C on solid Luria– Bertani (LB) broth medium with 100 μg/mL ampicillin. Colonies with confirmed presence of the correct insert by PCR were grown overnight in liquid LB with the appropriate antibiotics at 37°C and plasmids were isolated (GeneJET Plasmid Miniprep Kit, Thermo Fisher Scientific) and verified by sequencing. These constructs were used as template to fuse attb1 and 2 sites to the G4 sequence (without signal peptide). A 2-step PCR was performed with firstly gene-specific primers used in 20 μL of PCR mix with an annealing temperature of 45°C (five cycles), followed by 54°C (25 cycles). Secondly, PCR was performed using 3 μL from the previous mix in 17 μL of fresh PCR mix employing the primers attb1 short and attb2 short using the same program as described above. Obtained PCR products were purified as described above and ligated in the Gateway® pDONR™221 vector (Thermo Fisher Scientific Waltham, Massachusetts, USA). Transformed cells were grown o/n at 37°C on solid Luria–Bertani (LB) broth medium supplemented with 50 μg/mL Kanamycin and plasmids were isolated as described above.

Effector G2 and G1-diet (G1-d) without their native signal peptide, and flanked by attLR1 and attLR2 cloning sites were synthesized at GeneUniversal (https://www.geneuniversal.com/, Newark) in vector pUC57. Coding sequences of G4 interacting proteins MIPDB141 (Meloidogyne-induced giant cell protein DB141; Solyc01g099770.2) and RSP1 (REF-like stress related protein 1; Solyc05g015390.2.1) were retrieved from the Sol Genomics Network (SGN) database (Fernandez-Pozo *et al*., 2015), flanked with attLR1 and attLR2 cloning sites and synthesized at GeneUniversal in vector pUC57. In addition, the sequence part of MIPDB141 that was overlapping among the different partial cDNA clones (preys) interacting with G4 in the Y2H screen was synthesized as well in the same vector (Figure S1, see Supporting Information). The design of the VIGS construct was carried out by applying the full coding sequence in the Sol genomics VIGS tool (https://vigs.solgenomics.net/) and by using database *Solanum lycopersicum* ITAG v2.40 to select the best region to target for the gene silencing. The fragment sequences were synthesized at GeneUniversal as described above. The VIGS construct was covering part of mRNA of the genes Solyc01g099770 and Solyc01g099780 (Figure 1, see Supporting Information). For all ENTRY vectors ligation reactions (LR Clonase™II, Thermo Fisher Scientific Waltham, Massachusetts, USA) were performed to bring them in the desired destination vectors. Constructs were sequenced to exclude (frameshift) mutations before transformation of Agrobacterium. All information on effector and plant cDNA and protein sequences used in this study are presented in Figure S1 (see Supporting Information). Alignment with G1-t and G1-d is presented in Figure S2 (see Supporting Information). Information on plasmids used in this study are presented in Table S5 (see Supporting Information). Constructs were either transformed into Agrobacterium strain GV3101 using the freeze–thaw method described in (Holsters et al., 1978) for agroinfiltration in *N. benthamiana, N. tabacum* and VIGS in *S. lycopersicum* or in Agrobacterium strain 1D1249 for agroinfiltration *S. lycopersicum* and *C. annuum* for ROS and bioassays. Colonies with confirmed presence of the desired construct were grown for 2 days in liquid LB with the appropriate antibiotics at 28°C and 200 rpm.

**Figure 1:**
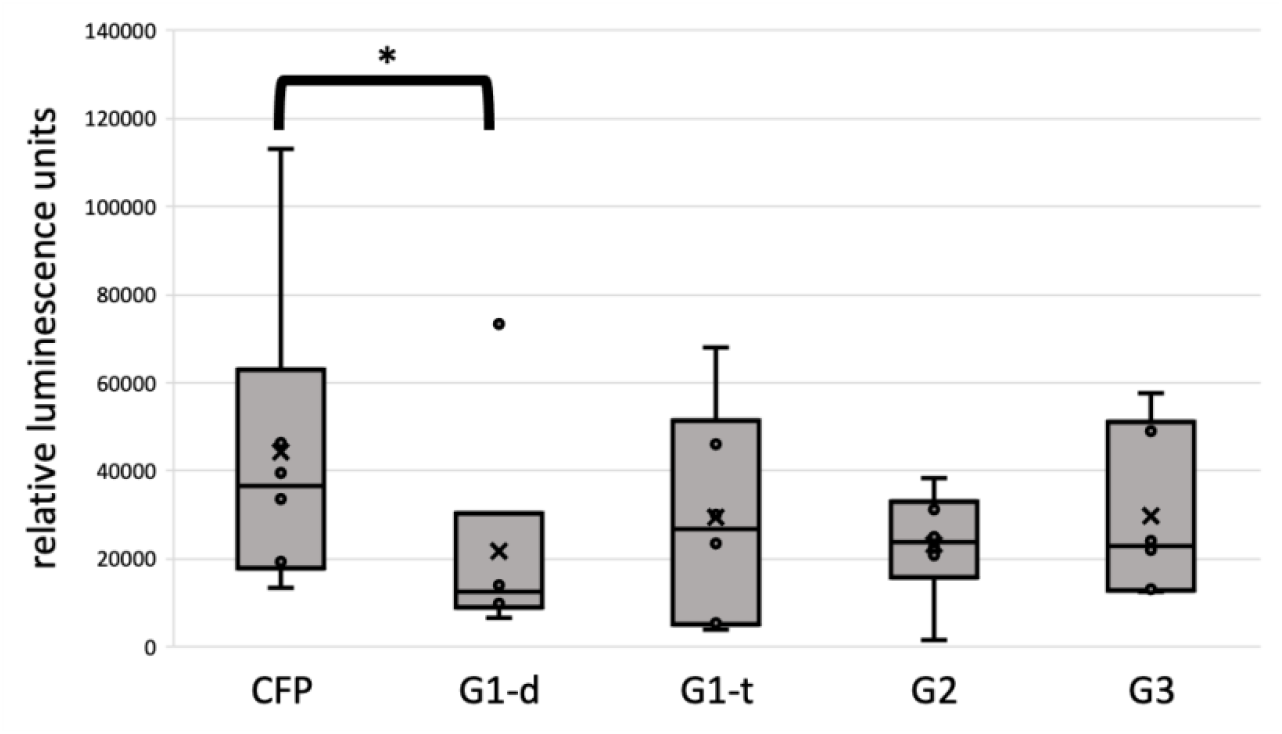
ROS assays with candidate whitefly effectors. Relative luminescence, representing reactive oxygen species (ROS) production during a flg22-induced assay in *Nicotiana benthamiana* transiently expressing candidate effectors G1, G2 and G3. CFP: cyan fluorescent protein as control. G1-d: G1 identified in artificial diet. G1-t: G1 identified in transcriptomic data. Asterisk (*P<0.05) indicate a significant difference compared to the CFP control shown by Friedman test (not normally distributed). Box plots represents RLU produced by 6 plants. Box shows the upper and lower quartiles, whiskers show the minimum and maximum data point within 1.5 x interquartile range. Line within box marks the median, cross marks the mean.

#### Modification of the destination vectors luciferase complementary assay

To carry out the split-luciferase complementary assay a red-shifted version of the firefly luciferase having the mutation I288A (Wang et al., 2013) was *Arabidopsis thaliana* codon optimized, synthetized (Eurofins genomics; for coding sequence Figure S1, see Supportive information) and cloned into destination vector pGWB402 (Nakamura et al., 2010). The N-terminally tagged luciferase and the empty vector PGWB402 were digested with the restriction enzyme *XbaI*. The C-terminally tagged luciferase and the empty vector PGWB402 were digested with the restriction enzyme *AfeI*. The fragments contain the luciferase coding sequence of amino acids 1-398 for N-terminus and 394-550 for C-terminus, which was previously found to be the combination with the lowest background signal (Paulmurugan and Gambhir, 2007). The N-terminus is fused in frame to a FLAG-tag and the C-terminus to a HA-tag. The fragments were PCR amplified and inserted in the gateway compatible binary vector pGWB402 by restriction cloning (Nakagawa et al., 2007). This eventually led to availability of the following vectors for in frame fusion of the protein of interest pGWB402 HA-RedFFLuc Cterm-Gateway box (attR1-attR2, standard frame, stop codon) and pGWB402 Gateway box - RedFFLuc Nterm-FLAG-Stop (no start codon) (Table S5, see Supporting Information).

### Assay for reactive oxygen species (ROS) induced by flg22

#### Infiltration of N. benthamiana

*Agrobacterium tumefaciens* (strain GV3101) carrying either effector constructs or, as negative control, the CFP construct was grown for 1-2 days at 28°C in LB medium supplemented with 25 μg/mL rifampicin, 50 μg/mL gentamicin and 50 μg/mL spectinomycin. The agrobacterium carrying the P19 silencer inhibitor construct (pBIN61:P19) (Jay et al., 2023) was grown 1-2 days at 28°C in LB medium with 25 μg/mL rifampicin and 50 μg/mL kanamycin. Liquid cultures with agrobacteria were centrifugated for 5 min at 3500 rpm (RT), resuspended in infiltration buffer (2% w/v sucrose, 1x Murashige & Skoog Basal Salt Mixture without vitamins (Duchefa, the Netherlands), 10 mM 2-(N-morpholino)-ethanesulfonic acid (MES), 200 μM acetosyringone, pH adjusted to 5.6) and diluted to an OD_600_ of 0.6. The cell solution was incubated o/n with slow shaking in the dark at RT, or when infiltration was performed at the same day, incubation was done for 3 h in the dark at RT without shaking. Prior infiltration, the Agrobacterium carrying the effector construct and Agrobacterium carrying the P19 silencing inhibitor were mixed in a ratio of 1:1. Agroinfiltration was performed by making a small hole in the abaxial site of the leave followed by infiltration using a needleless syringe. Plants were allowed for expression for circa 2 days.

#### *Infiltration of S. lycopersicum* and *C. annuum*

Agrobacterium (strain 1D1249) carrying either a effector construct or, as negative control, the CFP construct was grown for 1-2 days at 28°C in LB medium with 50 μg/mL spectinomycin. Liquid cultures with agrobacteria were centrifugated for 5 min at 3500 rpm (RT), resuspended in infiltration buffer and diluted to an OD_600_ of 0.3. Discs of 2,4 mm, including a small part of the petiole, from 4–5-week-old *S. lycopersicum* or *C. annuum* plants were taken from expanded leaves. Vacuum agroinfiltration was performed as described in (Abd-El-Haliem et al., 2018) with some minor modifications; Discs were placed in 6 wells plates with on the side 0,6 % Daishin agar dissolved in MQ water (Duchefa, the Netherlands). The bottom of the leave discs was placed in the agar and the plastic lid was placed back on the plate. Plates were placed on 25°C under 16h/8h light-dark regime in a climate cabinet for 3 days.

#### Luminescence measurement

Leaf discs of 16 mm^2^ were collected with a cork borer from the *N. benthamiana* infiltrated spots, or from the *S. lycopersicum* or *C. annuum discs* and transferred to 96-well white plates. Assays were performed for 6 to 8 plants and of each infiltration, two replicates were taken. The leaf discs were floated for 6 hrs-12 hrs on 190 μL autoclaved MQ water for recovery. Just prior the Luminescence measurement, the water was removed and replaced by a mixture of 100 nM flg22 (QRLSSGLRINSAKDDAAGLAIS; (Felix et al., 1999)), 0.5 mM luminol probe 8-amino-5-chloro-7-phenyl-pyrido[3,4-d]pyridazine-1,4(2H,3H)dione (L-012) (Wako Chemicals, Richmond, Virginia, USA) and 20 μg/mL horseradish peroxidase Type VI-A (Sigma, Saint Louis, Missouri, Verenigde Staten). ROS production was measured by a luminol-based assay (Keppler, 1989) over 45 min with measurement interval of 2 min with integration time of 1 sec.

#### Statistical analysis

Statistical analyses for ROS assays were performed by taking the average of the two discs from the same infiltration spot or discs. Total luminance of 8 plants was used for further statistical analysis. The data were analyzed for normality with SPSS and were either used in a student’s T test or Mann-Whitney U test. In case of more than two different effector constructs, either repeated measures ANOVA or Friedman test was used. Values of *P*≤ 0.05 were considered significantly different compared to the CFP control group.

### Whitefly bioassays

Agroinfiltration was performed as described for the ROS assays. After 3 days of expression in *N. tabacum*, a whitefly bioassay was performed as described in van Kleeff et al. (2022; under revision). Agroinfiltration on *S. lycopersicum* was performed as described under ROS assays. After 3 days, the leaf discs were placed in a Petri dish with vents, whereby to the edge of the Petri dish 0,6% agarose was poured and the solidified agar was covered with transparent foil (Figure S6A). The petiole was placed through a cut in the foil in the agar in such way that the leaf discs were sticking out from the side, leaving space for the whiteflies to move to the abaxial side of the leaf disc. White weighing paper was placed at the bottom of the Petri dish to reduce humidity in the dish and attachment of the whiteflies to the plastic. Petri dishes were closed and sealed using Leukopore tape. Whiteflies were collected from cucumber plants by aspiration in a glass Pasteur pipette sealed with parafilm and placed on ice until they reached a chilled-coma. 15 whitefly females were placed in each Petri dish and sealed with Leukopore tape. Petri dishes were placed in a growth cabinet at 27°C, 70% relative humidity and a 16h/8h light-dark regime. After 3 days the Petri dishes with whiteflies were stored at -20°C. Healthy looking females (undamaged and no morphological changes in color and hydration similar to whiteflies in chill-coma) were considered to be alive during freezing and counted as living females. Statistical analysis was performed on the eggs produced per living female using a similar approach as for the ROS assay.

### Yeast two-hybrid screening

A LEXA yeast two-hybrid screening was performed by Hybrigenics Services (France; http://www.hybrigenics-services.com) with the G4 coding sequence (excluding its native signal peptide) against a cDNA library of leaf, petiole, stem and root tissue of TYLCV-infected *S. lycopersicum* constructed into Activation Domain (AD) vector pP6 (TOPLI). G4 was PCR amplified using G4-pUC57 vector as template and cloned into a LexA DNA-Binding Domain (DBD) vector pB27 (N-LexA-G4-C fusion). Obtained prey fragments were sequenced and identified by the NCBI database. The Predicted Biological Score (PBS) following Formstecher et al. (2005) was used to indicate the confidence of an interaction. These scores led to four categories: very high confidence, high confidence, good confidence and moderate confidence. Selected preys were confirmed by 1-by-1 screening on selective medium without tryptophan and leucine (control) and selective medium without tryptophan, leucine and histidine (confirmation of interaction). As a negative control the construct were combined with empty pB27 vector or empty pP7 vector.

### Co-localization

*Agrobacterium* (strain GV3101) carrying either an effector construct or, as negative control, the RFP construct, were grown for 1 days at 28°C in LB medium supplemented with 25 μg/mL rifampicin, 50 μg/mL gentamicin and 50 μg/mL spectinomycin. Preparation of the liquid culture and agroinfiltration were performed as described above and the agrobacteria carrying the effector or plant protein fused to the fluorescence tag were infiltrated together with the P19 silencing inhibitor construct in infiltration buffer with an OD_600_ of 0.5, on the abaxial leaf side of 4-6 week old *N. benthamiana* plants. Expression was allowed for 3 days in 21°C under 16h/8h light-dark regime. For imaging, leaf discs were prepared from the infiltrated leaf area and placed on glass slides using double sided tape. Imaging of the fluorescence signal was performed on a Nikon Ti A1 confocal microscope using a 20x Plan Fluor, NA 0.75 (multi-immersion) objective. eRFP was excited with a wavelength of 561 nm and emission was detected at 592–632 nm. eCFP was excited with a wavelength of 440 nm and emission was detected at 465-500 nm nm. Auto-fluorescence of chlorophyll was detected at 657–737 nm.

### Luciferase complementation assay

Agrobacteria (strain GV3101) carrying the desired construct (vectors used can be found in Table S5, see Supporting Information) were grown o/n in liquid LB supplemented with 25 μg/mL rifampicin, 50 μg/mL gentamicin and 50 μg/mL spectinomycin. The liquid cultures were centrifuged and dissolved in infiltration buffer (as described in the section “*Assay for reactive oxygen species (ROS) induced by flg22*”) to an OD_600_ of 2. The complementary constructs were mixed in a ratio of 2:2:1 with the P19 silencing inhibitor, so a final OD_600_ 0.8 was reached for the Agrobacterium carrying the effector and plant protein constructs and a final of OD_600_ 0.4 was reached for the Agrobacterium carrying the P19 silencer inhibitor. Infiltration spots were circled on the leaves with a marker and numbered to identify the infiltrated constructs during luminescence measurement. Infiltration was performed using a needleless syringe on the abaxial side of the 4 till 6 weeks-old *N. benthamiana* plant leaves and each leaf was infiltrated in a cycle location to avoid location-specific signal. The agroinfiltrated *N. benthamiana* leaf tissue was examined 3 days post-infiltration. Leaves were detached from the plant and moved to Petri dishes with tissue saturated with MQ water. Leaves were brushed 2x times with D-luciferin buffer (2 μL D-luciferin/mL MQ (L1349, Duchefa Biochemie); 0,02 % SILWET ® L-77 (Kurt Obermeier GmbH& Co.KG; Bad Berleburg, Germany). After 2-4 hours in the dark the signal was detected using a charge-coupled device imaging system CCD camera (Princeton Instruments – 0302110003) to capture the chemiluminescence signals. Data acquisition was performed using the MetaVue program. Chemiluminescence was detected without any filters during an exposure of 5 minutes. For all observations, the binning was set at 10. For every experimental sample, eight independent biological replicates were examined. Raw pictures were analyzed and processed with ImageJ (https://imagej.nih.gov/ij/). The mean (integrated density/area size) with extraction of the leaf background signal was used for the statistical evaluation of the data. Normalization was performed by total signal of all samples. Normalized data of 8 plants was used for repeated measures ANOVA tests in SPSS. As a negative control effector G4 fused to N-terminus half of the luciferase complemented with TYLCV ^Alb13^ Rep (FJ956702.1) (Maio, 2019, dissertation), containing the C-terminus half of the luciferase was used.

### Co-immunoprecipitation

Co-immunoprecipitation was performed at Profacgen (https://www.profacgen.com/). For immunoprecipitation, constructs of interest were expressed in *N. tabacum* for two days and subsequently the leaf tissue was ground by liquid nitrogen. About 3 g tobacco powder was taken up in lysis buffer with protease inhibitors on ice for 15 min and the supernatant was collected by centrifugation. Protein A/G beads were washed three times with lysis buffer and empty Protein A/G beads were added to the supernatant for cleaning for 30 min. After cleaning, the Protein A/G beads were discarded, an appropriate amount of GFP antibody was added to the supernatant, and the beads are incubated again for 3 h. The cleaned Protein A/G beads are added again to fully bind the antibody for 2 h. After incubation, the beads were centrifuged at 800 g for 3 min at 4 °C and washed three times with 1×PBS. Loading buffer was added and placed in boiling water bath for 10 min and subsequently centrifuged for 10 min. Proteins were separated by SDS-PAGE, transmembrane blocked and incubated with primary antibodies (Anti-MYC and Anti-GFP 1:5000), secondary antibodies (Anti-mouse/Anti-rabbit (HRP, 1:5000)), and developed (ECL).

### Phloem exudate collection

Circa 4-week-old *S. lycopersicum* plants (Moneymaker) were placed in a large cage and exposed to a mixture of adult male and female whiteflies. Each week a new batch of whiteflies was added to the cage to get more variation in nymphal stages. After circa 4 weeks phloem extraction was performed on well-infested plant leaves. As negative control non-infested *S. lycopersicum* plants kept under similar environmental conditions were used. Phloem exudate from tomato was extracted as describe previously by Narvfiez-Vfisquez et al. (1994) and Madey et al. (2002) with slight modification. Briefly, four leaves from each tomato plants were excised at the petiole using scissors and immersed in bleeding buffer (5 mM phosphate buffer, 5 mM EDTA) for 20 min. The petioles were recut about 2 mm above original cut while immersed in bleeding buffer and transferred to 2 mL phloem collection buffer (5 mM phosphate buffer, 5mM EDTA and 0.5x protease inhibitor (Roche cOmplete EDTA-free Protease inhibitor)) and incubated for 6 h in humid conditions.

Proteins in the exudates were then concentrated by acetone precipitation (Mitton et al., 2009) and suspended in Ambic buffer (100 mM ammonium bicarbonate, 10 mM tris(2-carboxyethyl)phosphine, 40 mM chloroacetamide). All samples were quantified using Bradford assay and 25 μl of each sample was run in SDS-PAGE. Trypsin digestion was carried out by adding 1μg trypsin in 5 ng of total protein for overnight incubation at 37 ºC. Next day, 1% of formic acid was added to acidify the protein sample. For cleanup omix C18 column, omix C18 column was first activated by adding 100% acetonitrile, then washed three times using washing buffer (0.1 % formic acid). The sample was then added to column and washed three times using washing buffer. The sample then eluted using elution buffer containing 60% of acetonitrile, 0.1% formic acid and 39.9 % of water. 5 μl of sample then used for LC-MS analysis.

This analysis was performed as described in Tu *et al*. (2021). Raw MS/MS data were searched in Maxquant (version: 1.6.14.0) (Cox and Mann, 2008) against an in house created *B. tabaci* proteome database and *a S. lycopersicum* database downloaded from Uniprot (06/2020). To control for the false spectrum assignment rate (1% FDR), a reverse version of the same databases was also searched. Settings were set as default for timsDDA. Trypsin/P was selected as the digestion enzyme, with a maximum of 2 missed cleavages. The Oxidation (M) was set as a variable modification and Carbamidomethyl (C) as a fixed modification. For accurate mass, retention, and tims-time matching: “match between runs” was selected with a matching time window of 0.2 min and a matching ion mobility window of 0.05 indices.

### VIGS

#### Agroinfiltration

Agrobacterium carrying either the construct with the plant protein fragment, the GFP negative, PDS positive control (Liu et al., 2002) construct or the helper plasmid were grown o/n at 28°C in LB medium supplemented with 25 μg/mL rifampicin, 50 μg/mL gentamicin and 50 μg/mL spectinomycin while shaking at 220 rpm. The next day, 250 μL of the overnight culture was added to 20 mL LB medium supplemented with the same antibiotics, 10 mM MES and 20 μM acetosyringone and grown again o/n at 28°C and 220 rpm. The following day, the liquid cultures were centrifuged at RT for 5 minutes at 3600 rpm and the bacterial pellet was resuspended in infiltration buffer to a final OD_600_ of 0,5. Agrobacterium carrying either the protein-TRV2 constructs or the TRV1 construct were mixed at 1:1 (final concentration of each construct was 0.25). Finally, the mixture was incubated at RT at least 3 h in upright 50 mL tubes or o/n while slowly shaking while the tubes were laying down. Cotyledons of 8–9-day old *S. lycopersicum* plants (cv. Moneymaker) were completely infiltrated with the Agrobacterium mixtures carrying the desired construct and the helper TRV1 plasmid using a needleless syringe. After infiltration the plants were watered and grown in the conditions described under the topic plant growth until use for the whitefly bioassays 4-5 weeks post infiltration.

#### VIGS combined with whitefly bioassay

As a control for silencing effect, a batch of plants was infiltrated with a construct silencing the *PDS* gene. Based on the spread of morphological changes related to PDS silencing, leaves considered on having a change to be silenced for TCTPs were selected and detached from the plant. The leaf was used to cut a disc of 24 mm including a small part of the petiole. Then, the leaf disc was placed in a Petri dish as described in the section “*Whitefly bioassays*”). Plates were placed in a growth chamber at 25 °C, 70% relative humidity and a 16h/8h light-dark regime. The next day, whiteflies were collected from cucumber plants by aspiration and placed on ice until they reached a chill-coma. 10 whitefly females were placed in each Petri dish that was then sealed again with Leukopore tape. The plates were incubated in a growth chamber for 2 days at 27°C under a 16/8 h light-dark regime. After the incubation time the plates were placed at -20°C. Healthy looking females (as described in the section “*Whitefly bioassays*”) were considered to be alive during freezing and counted as living females. Statistical analysis was performed on the eggs produced per living female, using a factorial ANOVA.

#### VIGS combined with choice assay

Whitefly choice assays were performed on square Petri dishes (120 mm × 120 mm) whereby on opposite sides 0.6% plant ager was poured (while keeping the plates on the side) which was covered with transparent foil (Figure S6B). Small cuts in the foil were made to be able to place the discs in the agar, as described above. On one side 3 discs of the same *MIPDB141*-silenced plant were placed, while on the other side 3 discs of GFP silenced plants were placed. The three discs that were placed on one plate were taken from the same plant. In the middle of the plate 15 whiteflies in chill-coma were placed and the plates were sealed with Leukopore tape. The plates were placed in random orientation in the growth cabinet under the same conditions as described in the section “*VIGS whitefly bioassay*”. After 2 days, the plates were carefully transported to -20°C. After at least 3 h the plates were used for counting number of eggs and living females (considered to be alive prior to freezing). Scoring was performed by analysing the position of the whiteflies (GFP or MIPDB141 site). The number of eggs produced were divided by the number of females found on the corresponding site. Each bioassay was performed on 6 to 8 different plants for both the GFP control as the MIPDB141 silenced constructs. The choice assay was performed twice, with either 6 or 8 plates containing 15 whiteflies per plate. Statistical analysis was performed using a replicated G-test for Goodness of Fit.

## Results

### At least seven candidate effectors could be identified in artificial diet

To identify candidate effectors, adult whiteflies were fed with artificial diet that was analyzed on presence of proteins. Two *B. tabaci* proteins/protein groups, BTA026858.1 (S2, van Kleeff et al., 2022, under revision) and BTA009432.1 (S1, van Kleeff et al., 2022, under revision)/BTA009433.1, were identified in artificial diet collected after 6 hours of *B. tabaci* feeding. In artificial diet collected after 24 hours of *B. tabaci* feeding nine proteins/protein groups were identified BTA026858.1 (S2), BTA009432.1 (S1)/BTA009433.1 together with BTA002396.1 (G1-d), BTA023203.2 (G2), BTA007921.1 (G3), BTA018678.1, BTA003195.1/BTA027670.1, BTA007014.1/BTA027040.1 and BTA021638.1 (G4) (Table S2, see Supporting Information). For all samples, the highest number of peptides was found for the hypothetical protein BTA026858.1 (S2), and to a lesser extent for the hypothetical protein BTA009432.1 (S1). Based on a tblastn search, eight of the nine identified proteins (or protein groups) have been previously identified in *B. tabaci* (MED) artificial diet by Huang et al. 2021 (Table S3, Supporting Information). Based on another study, the genes encoding S1, S2 and BTA023203.2 (G2) are the three most highly expressed genes in *B. tabaci* MED heads, while the other identified proteins are encoded by genes belonging to the top 2% of most highly expressed genes in *B. tabaci* MED heads (Table S3, see Supporting Information (Wang et al., 2020)). Seven of the nine identified proteins/protein groups were predicted to include a signal peptide (Table S2). Eventually we continued with a subset of these proteins for further characterization using the following criteria: two proteins (groups) were excluded in further functional analysis as they were characterized in another project (S1 and S2) and further selection was made based on the presence of a signal peptide which led to the selection of four interesting candidate effector proteins (G1-G4). G1 was found to have a potential paralog gene (G1-t) in the *B. tabaci* MEAM1 transcriptome (Lee et al., 2018; van Kleeff et al., 2022, under revision), and therefore both were further analyzed for their role in immune suppression. Sequence information can be found in Figure S1 (see Supporting Information).

### *G4* encodes a protein with a predicted size of 267 amino acids and is whitefly specific

To obtain more insight in whether *G4* is present in *B. tabaci* MEAM1 and may be part of a gene family we performed a BLASTp search against the predicted proteome of *B. tabaci* MEAM1. This resulted in one identical hit (Bta08784) with G4 retrieved from *B. tabaci* MED protein database (BTA021638.1). The open reading frame of G4 contains 804 nucleotides (Figure S1, see Supporting Information) encoding a protein with a size of 267 amino acids, including a N-terminal signal peptide of 23 amino acids (likelihood 0,9569). A BLASTp search against the NCBI database (non-redundant proteins for all organisms) only resulted in hits with *B. tabaci* indicating this effector is specific for whiteflies. Using LOCALIZER, which specifically predicts where eukaryotic effector proteins will localize in plant cells (Sperschneider et al., 2017) the mature G4 protein was not predicted in the chloroplast, mitochondria or nucleus. However, WolfPsort, predicted a nuclear-cytoplasmic localisation for the G4 protein. Based on previously obtained *B. tabaci* transcriptomic data (Lee et al., 2018; van Kleeff et al. 2022, under revision) G4 is highly expressed in the salivary glands of adult whiteflies, while not being expressed in the nymphal stage. Last, G4 was potentially identified as a virulence effector using the VIVE assay, although this paper contains some uncertainties (Shi et al., 2020).

### G4 suppresses flg22-induced ROS in several plant species

To investigate whether the selected candidate effectors were able to supress the early immune response of the plant, we performed reactive oxygen species (ROS) inhibition assays in *N. benthamiana*. Effectors were transiently expressed *in planta* and leaf tissue was subsequently exposed to the PAMP peptide flg22. Expression of G1 resulted in a mixed response between performed ROS assays whereby in some cases significantly less H_2_O_2_ was measured compared to free CFP as negative control (Figure 1) and in other cases the H_2_O_2_ level was lower albeit not statistically significant. This was the case for both G1 retrieved from the artificial diet (G1-d) and the G1 paralog retrieved from the transcriptomic data (G1-t). Expression of G2 or G3 did not result in significant reduction in any of the performed ROS assays.

Effector G4 suppressed the flg22-induced ROS significantly compared to the CFP control (Figure 2) in *N. benthamiana* and was the most robust (3 out of 4 assays) suppressor of the 4 candidate effectors selected from the artificial diet. To test whether this effector may also affect immune responses of other plant species we performed the flg22 induced ROS assay also using leaf discs of the crop plants *S. lycopersicum* and *C. annuum* transiently expressing G4 (Figure 2). In both these plant species, G4 reduced the ROS production, but not as efficiently as observed in *N. benthamiana*. In the early stage (first 18 min, highest peak in CFP burst), G4 was significantly reducing the ROS production in *S. lycopersicum*, and caused a delay in the peak of the burst, but the total production of ROS after 45 min was almost similar to the free CFP control. In addition, G4 suppressed ROS in *C. annuum* in the first 23 min but the final ROS production was not significantly different after 45 min either. In contrast to *S. lycopersicum* the ROS burst in *C. annuum* was induced similar as in CFP control, but the ROS production was lower in the early interval of the burst. A representation of the ROS production for each time point can be found in Figure S3 (see Supporting Information).

**Figure 2:**
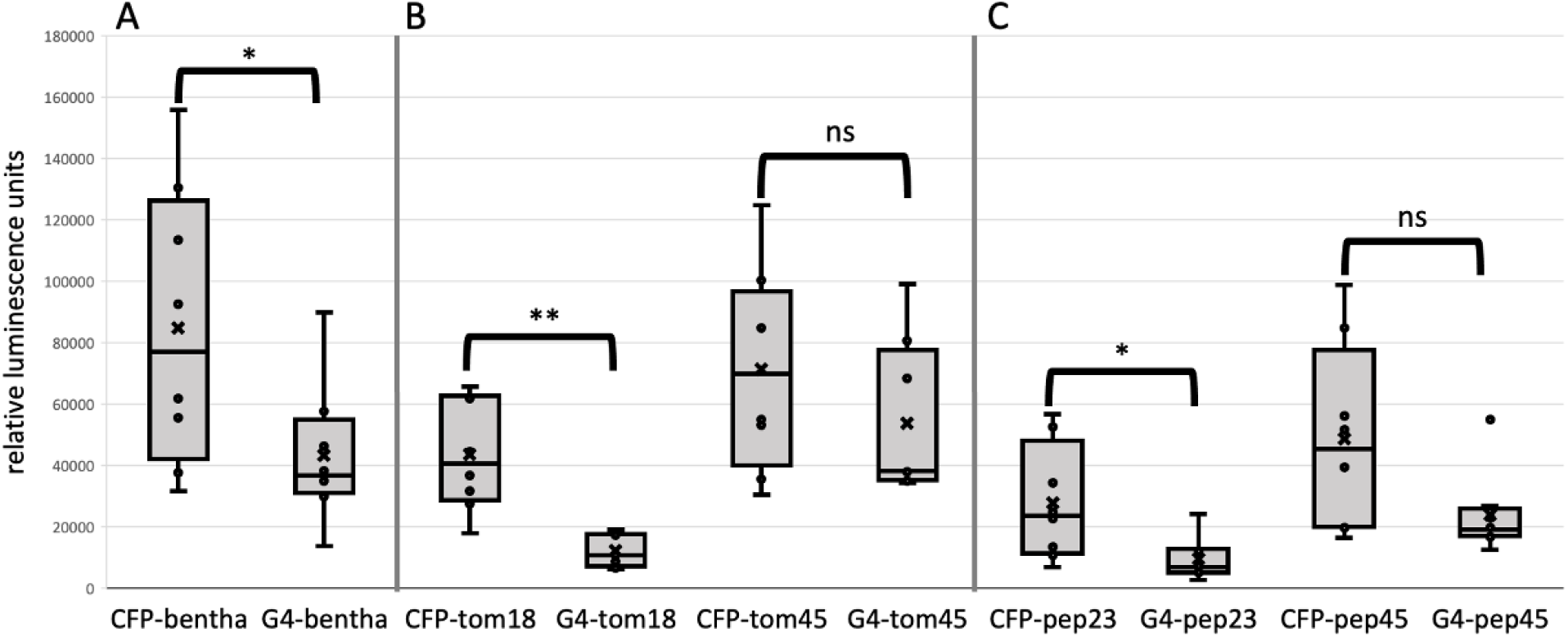
ROS assays with effector G4 in different plant species. Relative luminescence during a flg22 induced reactive oxygen species (ROS) assay in *Nicotiana benthamiana, Solanum lycopersicum* (Moneymaker) and *Capsicum annuum* transiently expressing G4. A: ROS production in *N. benthamiana* was analyzed 45 min after induction, showing a significant reduction in ROS production. B: ROS production in *S. lycopersicum* was analyzed 18 and 45 min after induction. C: ROS production in *C. annuum* was analyzed after 23- and 45-min. CFP: cyan fluorescent protein as control. Asterisks (*P<0.05, **P<0.01) indicate a significant difference compared to the CFP control shown by paired T-test or Mann Whitney-U-test when the data was not normally distributed. ns indicates non-significant difference compared to CFP control. Box plots represents RLU produced by 8 plants. Box shows the upper and lower quartiles, whiskers show the minimum and maximum data point within 1.5 x interquartile range. Line within box marks the median, cross marks the mean.

### Yeast two-hybrid screen identified two distinct *S. lycopersicum* target proteins of G4

To analyse whether G4 can directly interact with proteins in tomato, a LexA yeast two-hybrid screen was performed with G4 (excluding its native signal peptide) as bait, in yeast transformed with a cDNA library of leaf, petiole, stem and root tissue of Tomato yellow leaf curl virus (TYLCV)-infected *S. lycopersicum* (Hybrigenics Services, Paris, France, hybrigenics-services.com). A total of 94.1 million clones were screened and subsequently 253 His+ colonies were selected on a medium lacking tryptophan, leucine and histidine. This resulted in 52 primary candidates. A selection of the candidate target proteins was made, based on (i) (very) high confidence of interaction in the Y2H screen, (ii) putative role in immune suppression/stress response as described in literature, (iii) their presence in phloem exudates, or because its homolog in other plant species was a target of pathogen effectors that have been previously published. This led to a list of 6 candidate targets; Solyc01g099770.2 (MIPDB141); Solyc05g015390.2.1 (RSP1); Solyc10g045380.1.1 (Vacuolar protein sorting protein 62); Solyc02g067390.2.1 (RNA recognition motif containing protein); Solyc07g006280.2.1 (Senescence-associated protein; Tetraspanin) and Solyc08g074290.2.1 (BRI1-KD interacting protein 129). These interactions between obtained preys and effector G4 were confirmed by a 1-by-1 Y2H-interaction assay (Figure S4, see Supporting Information; Hybrigenics Services, Paris, France, hybrigenics-services.com).

### Proteomics of phloem exudates confirmed the presence of MIPDB141 in tomato

To determine whether these target proteins were present in the phloem, which support the probability of interaction in the host, and to analyse whether whiteflies affect the presence of these proteins, phloem exudates were collected from *S. lycopersicum* leaves infested with whiteflies and clean control leaves. MIPDB141 was detected in phloem of both whitefly-infested as well as non-whitefly infested plants (Table S4, see Supporting Information). In the phloem of non-infested leaves, TCTP was also found (Table S4, see Supporting Information), while this was not the case in the infested leaves. RSP1 was not detected in any of the phloem samples. In addition, we analyzed whether the candidate effector proteins identified in the artificial diet were detected in the phloem and although many other whitefly proteins could be detected in the phloem extract (data not shown), none of these proteins could be linked to the salivary proteins described in this study.

### Luciferase complementation assay confirmed interaction of G4 with both targets

To determine whether the interaction between G4 and the six selected targets could also be observed *in planta*, a luciferase complementation assay was performed with full-length plant proteins in *N. benthamiana*. Only RSP1 and MIPDB141 eventually resulted in a significantly increased luciferase signal (Figure 3). In addition, we aligned the amino acid sequences encoded by the partial cDNAs of MIPDB141 shown to interact with G4 in the yeast two-hybrid screen. The overlapping region was also cloned and expressed for analysing the interaction with G4 using the luciferase assay (This part of sequence is indicated in Figure S1 and S2, see supporting information). This was done to determine whether this region in MIPD141 interacts with G4. A significantly increased signal was observed with this part of the MIPDB141 protein confirming that an interacting site is located in this region. In addition, an increase in signal was also observed when G4 fused to both luciferin complements and co-infiltrated, showing G4 interacts with itself as well (Figure 3).

**Figure 3:**
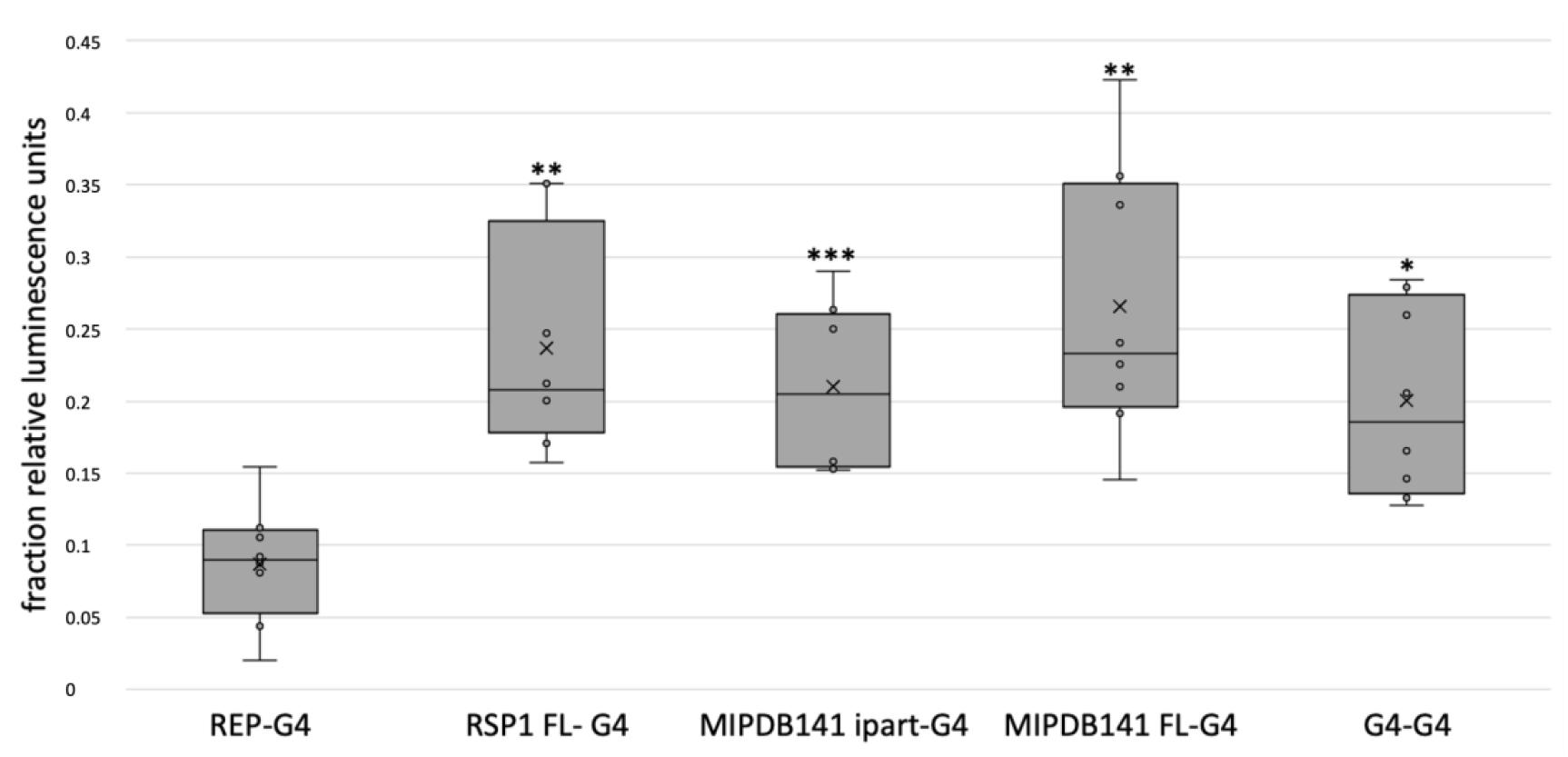
Confirmation of interaction of G4 with target plant proteins RSP1 (Solyc05g015390: REF-like related stress protein 1) and MIPDB141 (Solyc01g099770; Meloidogyne-induced giant cell protein DB141) using a luciferase complementation assay. REP-G4: negative control, FL: full-length coding sequence, ipart: interacting part (Figure S1). Asterisks (*P<0.05, **P<0.01, ***P<0.001) indicate a significant difference compared to the negative control shown by repeated measures ANOVA. Box plots represents the fraction of RLU produced by 8 plants. Box shows the upper and lower quartiles, whiskers show the minimum and maximum data point within 1.5 x interquartile range. Line within box marks the median, a cross indicates the mean.

### Co-immunoprecipitation confirmed the interaction between G4 and RSP1

To further confirm the interaction *in planta* between G4 and MIPDB141 and between G4 and RSP1 a Co-immunoprecipitation (Co-IP) was performed. All expressed proteins were detected in the input samples prior to GFP immunoprecipitation (GFP-IP), using tag-specific antibodies (Figure 4). MIPDB141 was not detected in the GFF-IP in both combination with free GFP (negative control) and G4 fused to GFP, indicating the interaction could not be further established with this assay (Figure 4A). RSP1 was only detected in the GFP-IP when GFP was fused with G4, with the faint signal indicating a weak interaction between G4 and RSP1.

**Figure 4:**
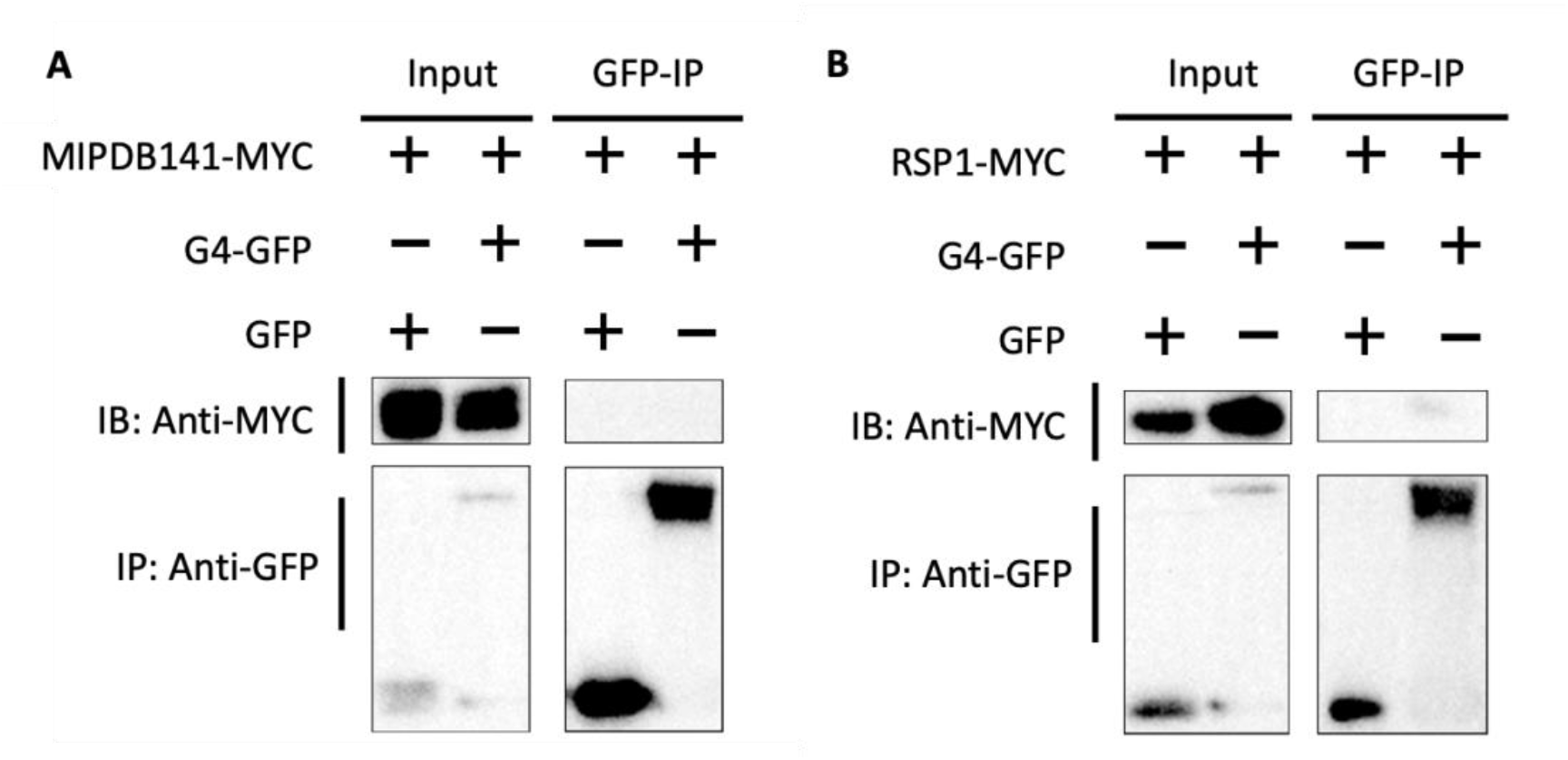
Co-immunoprecipitation G4 with MIPDB141 (Solyc01g099770; Meloidogyne-induced giant cell protein DB141) and target plant proteins RSP1 (Solyc05g015390: Ref-like related stress protein 1) in *Nicotiana tabacum*. A: MIPDB141 fused to Myc-tag was co-expressed with either G4 fused to GFP or free GFP. Proteins were expressed and detected in the protein extract. Either with free GFP or G4 fused to GFP, MIPDB141 was not detected in the GFP-IP. B: RSP1 fused to Myc-tag was co-expressed with either G4 fused to GFP or free GFP. Proteins were expressed and detected in the protein extract. RSP1 was detected in the GFP-IP when co-expressed with G4fused to GFP, but not with free GFP. IP: immunoprecipitate; IB: immunoblot.

### Subcellular localization and co-localization

To obtain insight on where G4 and the target proteins MIPDB141 and RSP1 accumulate in the plant cell, a subcellular localization of G4 fused either C- or N-terminally to CFP or RFP and MIPDB141 and RSP1 fused C- or N-terminally to RFP was performed when transiently expressed in *N. benthamiana* leaves. With all 4 constructs, the localization of G4 was observed in the cytoplasm and was excluded from the nucleus (Figure 5). G4 was also observed surrounding the nucleus, possibly accumulating in the ER. The proteins that could be confirmed as interactors *in planta* using the luciferase complementation assay were further analyzed for their subcellular localization as well. The localization of the candidate target protein RSP1 was determined in the cytoplasm with exclusion from the nucleus. MIPDB141, showed accumulation in the cytoplasm and nucleus, excluding the nucleolus (Figure 5).

**Figure 5:**
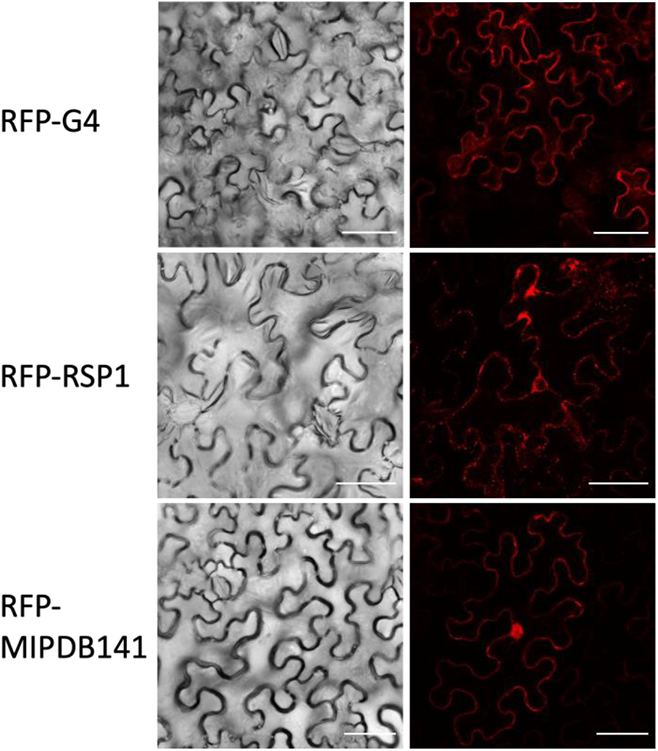
Subcellular localization of G4, RSP1 and MIPDB141 fused to RFP N-terminally, transiently expressed in *N. benthamiana*. G4 was observed in the cytoplasm and surrounding the nucleus, possibly accumulating in the ER. RSP1 was present in the cytoplasm and MIPDB141 in the cytoplasm and nucleus, excluding the nucleolus. Scale bar = 50 μm.

To determine whether the candidate targets were localized in the same cell compartments a co-localization assay was performed (Figure 6). Co-expression of RSP1 fused C- or N-terminally to RFP somehow resulted in lower protein accumulation of G4 fused to CFP C-terminally (data not shown). Both RFP orientations with RSP1 showed clearly co-localization with G4 fused to CFP C-terminally in the cytoplasm. During colocalization studies of MIPDB141 fused to RFP C- or N-terminally with G4 fused to CFP N-terminally, the latter was not detectable (data not shown). MIPDB141 colocalized with G4 in the cytoplasm, but not in the nucleus. There was no indication that G4 may play a role in inhibiting movement of MIPDB141 into the nucleus. None of the target proteins showed a different localization as result of co-expression with G4.

**Figure 6:**
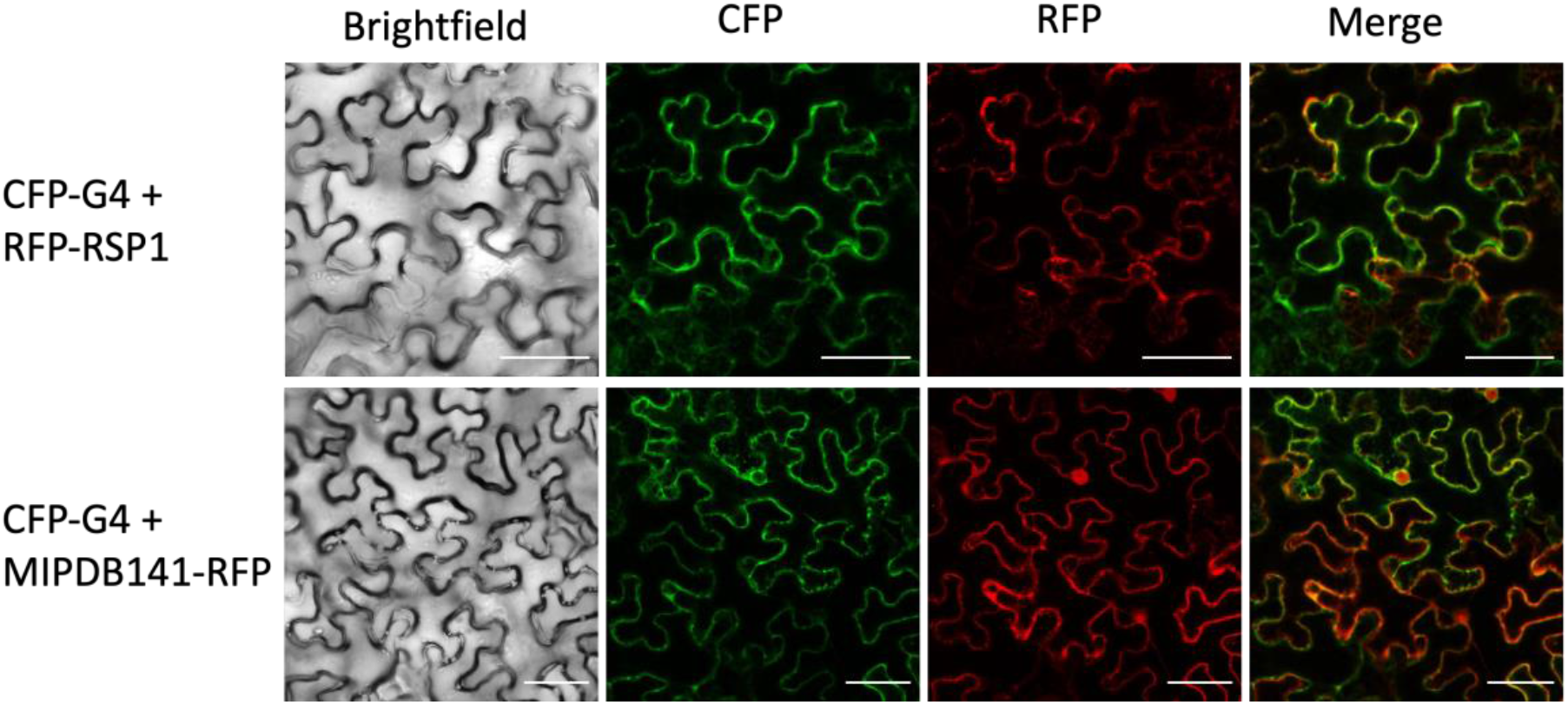
Subcellular colocalization of G4 fused CFP C-terminally and target proteins RSP1 and MIPDB141 fused to RFP C- and N-terminally respectively, after being transiently expressed in *Nicotiana benthamiana* leaves. RSP1 co-localized with G4 in the cytoplasm. MIPDB141 co-localized with G4 in the cytoplasm, but not in the nucleus. Scale bar = 50 μm.

### *In planta* expression of *G4* did not result in an altered whitefly performance

To determine whether excessive presence of G4 *in planta* would influence whitefly performance, we transiently expressed G4 in *N. tabacum* and *S. lycopersicum* and analyzed oviposition over 3 days. In both plant species the oviposition of the whiteflies was not significantly different compared to the CFP control group (Figure S5).

### Virus induced gene silencing of *MIPDB141* affects whitefly performance

To determine whether the target gene MIPDB141 plays a significant role in the whitefly-plant interaction we reduced its expression *S. lycopersicum* and assessed whether this affected whitefly performance. Concomitantly the MIPDB141 paralog (Solyc01g099780.2.1; TCTP) was also targeted by the VIGS constructs as it was impossible to design a paralog selective construct. We selected *MIPDB141/TCTP*-silenced leaves based on the white morphology of *PDS* silencing in control plants. Discs were taken and infested with female whiteflies for 2 days to allow oviposition. After 2 days of oviposition, the number of eggs per living female was significantly lower on the *MIPDB141/TCTP*-silenced plants compared to the GFP control (Figure 7A). Since a reduced oviposition was observed on the *MIPDB141/TCTP*-silenced leaf discs, a bioassay combining silenced and control leaf discs was performed to determine if reduced oviposition could be explained by a lower attractiveness of the *MIPDB141/TCTP*-silenced leaf discs relative to the controls for whiteflies. Unexpectedly, we observed that adult females were significantly more attracted to the *MIPDB141/TCTP*-silenced leaf discs compared to the GFP control (Figure 7B).

**Figure 7:**
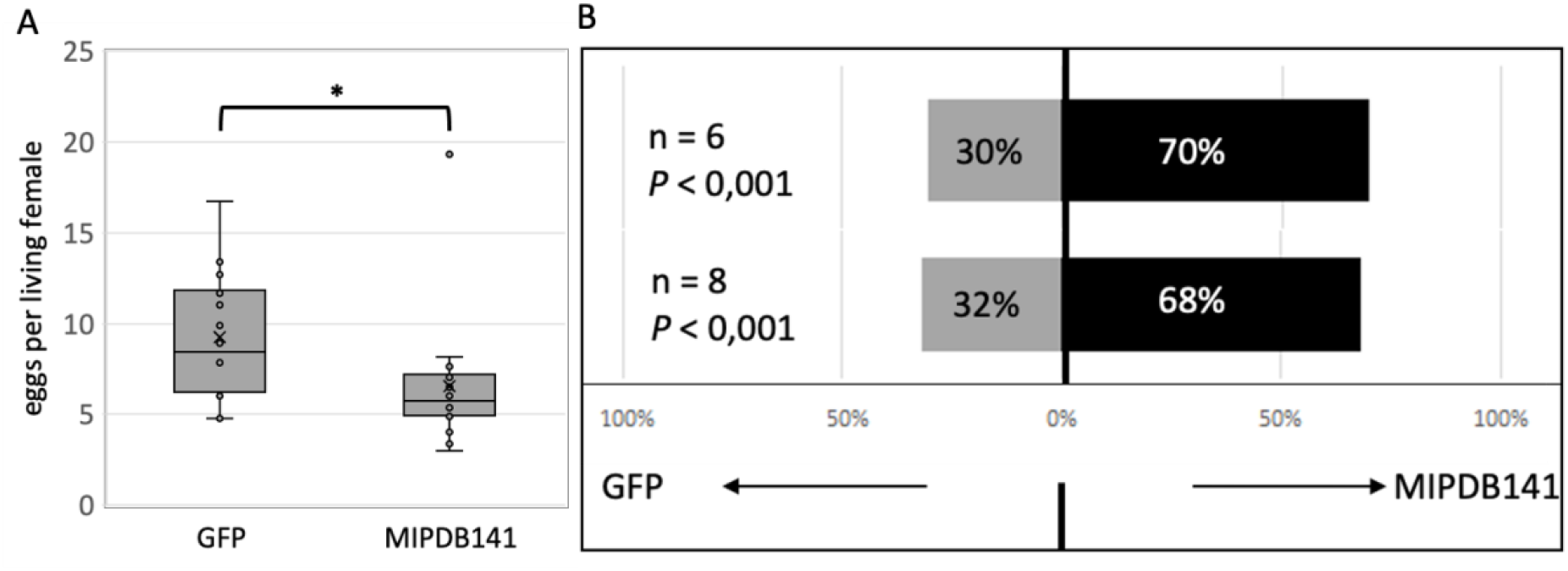
VIGS whitefly bioassays on *S. lycopersicum* (Moneymaker) with dsRNA produced against GFP (negative control) and MIPDB141. A: Number of eggs deposited by whiteflies in a non-choice assay after 2 days of oviposition (n = 16). Box plots represents eggs per living female on 16 plants. Box shows the upper and lower quartiles, whiskers show the minimum and maximum data point within 1.5 x interquartile range. Line within box marks the median, cross marks the mean. Asterisk indicate a significant difference compared to the GFP control shown factorial ANOVA (P < 0.05). B: Percentage of whiteflies that selected the GFP-silenced plants or the MIPDB141-silenced plants in a choice assay conducted twice independently. To determine significant difference a replicated G-test for Goodness of Fit was used.

## Discussion

Here we have shown that the effector protein G4 impairs tomato immunity to whiteflies by interfering with the elicitor-induced ROS response and via a direct interaction with tomato susceptibility protein MIPDB141. The identification and characterization of effectors used by herbivorous pests to manipulate the physiology of their plant host is an essential element for understanding the mechanisms that drive the ecology and evolution of plant-insect interactions. Moreover, elucidation of effector targets in host plants offers novel opportunities for resistance breeding to improve crop resilience, for example via knockout or mutagenesis of targets (Bisht et al., 2019; Vleeshouwers and Oliver, 2014). Different methods were successfully used to identify effectors of insect pests like *B. tabaci* (reviewed in Naalden et al. (2021)). In this study we used artificial diet as a food source for whiteflies as a direct approach for harvesting salivary proteins secreted by the inset via their stylets into their diet while feeding. Previously, this was successfully applied for identifying salivary proteins of spider mites (Jonckheere et al., 2016) and several insect species, including whiteflies (Huang et al., 2021; Yang et al., 2017). We were able to identify several proteins that qualify as candidate salivary effectors of *B. tabaci*. Of particular note, these proteins were previously also detected in artificial diet by Huang and co-workers (2021), indicating a robust secretion of these proteins by whiteflies.

To assess if our candidate effectors are present in the phloem of whitefly infested plants, we collected phloem exudates of whitefly-infested *S. lycopersicum* leaves, but none of the candidates here reported could be detected. Possibly some whitefly salivary proteins might have moved to, or even beyond, phloem companion cells as for example is seen with phytoplasma effectors (reviewed in Jiang et al. (2019)) or have a relatively short half-life after injection into the plant. Alternatively, the effector proteins identified from artificial diet might be specifically expressed by adult whiteflies, while during collection of the phloem exudates, taking several hours, adult whiteflies had been removed and only nymphs and eggs were present. Accordingly, van Kleeff and co-workers (2022, under revision) could identify only one whitefly effector candidate from phloem exudates and its corresponding gene was specifically expressed in nymphs and eggs. Hence, a different setup that does not exclude feeding adults during phloem extraction may give a better view on the whitefly salivary proteins injected during feeding.

Plants defend themselves against herbivores in various ways. The production of ROS is a central defence-related response in plants under attack as ROS are important defence signals as well as toxic for many insects (Goggin and Fischer, 2022; Kerchev et al., 2012). Sugarcane aphids for example, induces ROS production in sorghum and an H_2_O_2_ accumulation directly increases plant resistance against the aphid in different sorghum genotypes (Pant and Huang, 2021). In addition, feeding by whitefly nymphs resulted in upregulation of several genes in *A. thaliana* involved in scavenging of ROS and redox homeostasis, suggesting that nymphs induce ROS accumulation (Kempema et al., 2007). We performed a ROS induction assay to identify effectors that can suppress it. G4 acted as ROS suppressor, indicating that this protein may play a role as effector to modulate the immune response of the host during the whitefly-plant interaction. Due to the clear ROS suppression by G4 in *N. benthamiana*, we continued analysing ROS suppression in *S. lycopersicum* and *C. annuum* to confirm the effector is an important tool to search for susceptible genes. G4 gave consistently the most robust result in suppressing ROS in *N. benthamiana*, while in *S. lycopersicum* it delayed the ROS accumulation dynamics. These different effects on the ROS accumulation dynamics of different plant species could have many causes, such as differences in interaction strength between G4 and target orthologs, but on the other hands may just be a direct result of lower expression in *S. lycopersicum*.

Two target proteins of G4 in *S. lycopersicum* were identified in this study: RSP1 and MIPDB141. The interactors were identified by using a tomato cDNA library in yeast but were subsequently confirmed independently via *in planta* interaction assays. Concomittantly, the biological relevance of the G4: MIPDB141 interaction was confirmed via VIGS. Functional interactions of pathogen effectors with multiple host targets have been reported before (Pennington et al., 2016). Also, the whitefly orthologous effectors Bt56 and Bsp9 from *B. tabaci*, respectively, MED and MEAM1, were found to interact with two distinct plant proteins but it was confirmed in yeast that Bsp9 interacts also with the Bt56 plant target KNOTTED 1-like homeobox transcription factor (NTH202) supporting that whitefly effectors may have multiple targets in the host plant (Wang et al., 2019; Xu et al., 2019). We confirmed the interactions between G4 and the two plant targets *in planta* independently using the luciferase complementation assay. This is a relatively easy technique to screen interactions in plant tissue, and confirms interacting proteins are accumulating in the same cell compartment(s). Furthermore, using co-immunoprecipitation (Co-IP) as a fourth independent interaction assay, we could confirm the interaction between RSP1 and G4. Co-IP is especially suitable for relatively strong interactions (Struk et al., 2019). However, the interaction between MIPDB141 and G4 is probably relatively weak since it was not observed in the CoIP but could still be confirmed in the yeast and luciferase complementation assay, techniques that are both also suitable for relatively weak interactions. Altogether, we could confirm RSP1 and MIPDB141 to be *in planta* target proteins of G4. In addition, for MIPDB141 we also could confirm the interaction region luciferase complementation assay by using the fragment that is overlapping in the different truncated preys from the Y2H screen (Figure S1 and S2). Modification in this region might lead to a disturbed interaction between G4 and MIPDB141 which might be interesting for further resistance breeding.

MIPDB141, also known under the name Translationally Controlled Tumor Protein (TCTP) in other plant species, is highly conserved and has been identified in all eukaryotic organisms (Bommer and Thiele, 2004). It is involved in many cellular processes, like growth and development and (a)biotic stress responses (reviewed in Deng et al., (2016)). Transcriptome analysis of root-knot nematode-induced giant cells showed an upregulation of *MIPDB141*, indicating a role in nematode-plant interaction (Bird and Wilson, 1994). We were able to detect MIPDB141 in the phloem exudates of *S. lycopersicum*. TCTPs have been detected in phloem sap also in previous studies in different plant species like for example *Cucurbita maxima* (Aoki et al., 2005) and *A. thaliana* (Berkowitz et al., 2009). Since whiteflies in general are believed to secrete most of their effectors into the sieve tubes of the phloem, this provides strong support that a MIPDB141-G4 interaction *in planta* might indeed occur in the vascular bundle and/or its companion cells. Interestingly, Deng and co-workers (2016) showed that the MIPDB141 ortholog of *Hevea brasiliensis* (HbTCTP1) interacts with HbRSP1, its homolog of SlRSP1. Whether SlMIPDB141 and SlRSP1 also interact in *S. lycopersicum* is unknown, but given the high degree of conserveness of these proteins it is likely that this interaction occurs as well. Since G4 interacts with both MIPDB141 and RSP1 - and with itself - the three proteins may form a complex together and possibly G4 stabilizes the interaction between MIPDB141 and RSP1. Alternatively, G4 may destabilize the interaction between MIPDB141 and RSP1. Although we did not observe a change in localization when MIPDB141 was co-expressed with G4, co-expression with RSP1 in the same tissue too may alter localization if the two target proteins can form a complex and if G4 can influence it. The fact that we did not detect RSP1 in the phloem exudates, may be explained by accumulation of RSP1 in the companion cells to where effectors may also be translocated after secretion.

(Co-) localization studies shows that G4 and the two target proteins all localize to the cytosol but that MIPDB141 also localizes to the nucleus. The interaction of the two target proteins with G4 takes place exclusively in the cytosol and G4 does not influence localization of MIPDB141 to the nucleus nor does it follow MIPDB141. Both G4 and RSP1 did not localize in the nucleus, although both proteins fused to the fluorescence tag are sufficiently small to diffuse to the nucleus (Wang and Brattain, 2007). Whether the localization of G4 and RSP1 is either in the sieve tubes or in the companion cells could be further investigated using immunolocalization. For example, immunolocalization on the whitefly effector BtFer1 was performed by (Su et al., 2019) showing that it localizes to the phloem.

REF-like proteins in *H. brasiliensis*, orthologs of tomato RSP1, are stress-related proteins (Ko et al., 2003) which indicates that G4 might be involved in modulating the stress response in the plant tissue. TCTPs are highly conserved and many functions in the cell are attributed to these proteins, but there are relatively few *in planta* studies compared to studies in animal tissues or cell lines (Deng et al., 2016). This makes it challenging to predict the effect of G4-binding to RSP1 in plant tissues. HbTCTP1 in *H. brasiliensis* was shown to be regulated under several stress conditions including drought, wounding and H_2_O_2_ treatment (Deng et al., 2016) while overexpression of *AtTCTP* in *N. benthamiana* resulted in a decrease in cell death (Hoepflinger et al., 2013). Plants in general contain one to three different TCTP paralogs (Gutiérrez-Galeano et al., 2014). In tomato two TCTPs are present: MIPDB141 and Solyc01g099780.2. In our VIGS assay we most likely interfered with transcript accumulation of both paralogs possibly preventing functional redundancy that could potentially undo the effect of silencing. In the Y2H screen the main prey found was MIPDB141, although one of the positive colonies contained Solyc01g099780.2. The fact that the paralog was found only once may be simply due to underrepresentation of this transcript in the (unnormalized) cDNA library that had been used for the Y2H screen. It strongly suggests that G4 can interact with the two paralogs in tomato simultaneously since both are expressed in the same tissues under similar conditions (Tomato Expression Atlas; https://tea.solgenomics.net/). Taken together, the effect we observed in the VIGS assay can therefore very well be due to simultaneous silencing of both genes. In future research, it would be interesting to further explore whether both targets are equally important and whether mutations in both TCTP genes or in just *MIPDB141* could lead to an increased resistance against whiteflies.

The choice assay showed that whiteflies preferred the *MIPDB141*-silenced leaf discs. Yet, during the no-choice bioassay on *MIPDB141*-silenced leaf discs their oviposition was reduced relative to the control group confirming the insect needs a functional MIPDB141 for optimal performance. This points to a puzzling role for MIPDB141 in interacting with the whitefly and suggests that its foraging behaviour is influenced in the opposite way as its oviposition behaviour. At least we show here that the reduced number of eggs found on the *MIPDB141*-silenced leaf discs is not a direct result of whiteflies repelled by this plant tissue. Assuming the G4-MIPDB141 interaction is adaptive to the whitefly it may under natural circumstances never encounter plants without *MIPDB141* expression. Therefore, silencing this gene may generate irrelevant pleiotropic cues that influence foraging or oviposition behaviour in an unnatural manner. So from a biological point of view this result may not be very relevant. More important is that the whitefly needs a functional MIPDB141 protein for maximal performance. For breeding purposes, simply knocking the gene out may not be suitable for obtaining resistant crops because this conserved gene has several primary functions in the plant (Berkowitz et al., 2009; Hoepflinger et al., 2013) while, according to our data, it may make the plant more attractive to whiteflies, possibly negating the beneficial effect via oviposition reduction. Thus, the challenge will be to identify alternative MIPDB141 alleles, either from natural (resistant) plant populations or via mutagenesis, that encode for MIPDB141 proteins that do not interact with G4, thereby reducing whitefly performance, retain their primary function. In addition, these plants most likely will not attract whiteflies more strongly. Stacking such alleles in crops could be very valuable for breeding programs that aim for sustainable insect resistance.

## Supporting information

Supportive Information

## Contribution of authors

DN, RS, MRK, TVL and JV conceived the experiments. DN performed the cloning procedures, ROS assays, VIGS and bioassays. MM performed the co-localization. AI collected artificial diet with whitefly saliva, GB performed LC-MS/MS experiments and WD analyzed LC-MS/MS data of artificial diet. SD performed phloem extraction, GK and WR performed LC-MS/MS experiments and analyzed LC-MS/MS data of phloem exudate. NG and MK performed the vector adaptation for split-luciferase and wrote material and methods for this part. HVDB supervised NG and MK. DN, WD, MRK and RS wrote the manuscript. All authors have read and agreed on the content of the manuscript.

## Acknowledgement

The authors would like to thank the Van Leeuwenhoek Center for Advanced Microscopy. (LCAM; University of Amsterdam) for use of microscopes, Rodrigo Therezan de Freitas for guidance in the VIGS assay and Harold Lemereis and Ludek Tikovsky for the excellent plant care. This research is funded by the European Union’s Horizon 2020 research and innovation program (773 902-SuperPests) and by the Dutch Research Council (NWO) grant 19391 (TTI-VICI). SD was supported under the European Union’s Horizon 2020 research and innovation program (817526—PRE-HLB). The Topsector T&U program Better Plants for Demands (grant 1409-036 to HB), including the partnering breeding companies, supported the work on the vector optimalization related to the luciferase complementation assay.

