## Supplementary material for "Whitefly effector G4 interacts with tomato proteins of which MIPDB141 affects whitefly performance": Supportive Information

### Supporting Information

>G1-artificial-diet_Bta13784_BTA002396.1 cDNA
ATGCACGTGTACTTTATCTCAGCGGTAGCTGTCAGTATCTTTTCAAACTTTCAGGAAGCG
GATGGTTGGAAGGCGACAATTTATGATGATGTACAACATGGCAAGACTTTGGAAACTTTG
TCGGGAGGCCCTTGCCAGGAGCCGAAAAGCTACAATAACGACAAAGCTTCCTCAATCAAC
ACTTGGGGTGGGTGCGTGGTAATTTATAGTGATTACATGTGCAAAGGCACTAACAAAACG
ATGAAACCCGGGAGTCCTCACCATAAAGATTTCAAACTGCTACACTTCGATAATGTGCTA
TCTTCCATCGGTCCCTGCCCATAA

>G1-artificial-diet_Bta13784_BTA002396.1 Protein
MHVYFISAVAVSIFSNFQEADGWKATIYDDVQHGKTLETLSGGPCQEPKSYNNDKASSIN
TWGGCVVIYSDYMCKGTNKTMKPGSPHHKDFKLLHFDNVLSSIGPCP

>G1-transcriptome_salivary_glands_Bta14287 cDNA
ATGCACGTGTACTTACTCTCAGCGGTCGCTATCAGTATCTTTTTAAACTTTCAGGAAGCG
GTTGGTTGGAAGGCGACAATTTATGATGATGTACAACATGGCAAGACTTTGCAAACTTTG
TCGGGAGGCCCTTGCCAGGAGGTGAAAGACGCCAATAATGACAGAGCTTCCTCAATCAAC
ACTTGGGGTGGGTGCGTGGTTATTTATAGTGATTTTCAGTGCAAAGGCACCAACAAAACG
ATGAGACCCGGGAGTCCTCATCATAAAGATTTCAAACTGCTACACTTCGATAATGTGGTA
TCTTCCATCGGTCCTTGCCCTTAG

>G1-transcriptome_salivary_glands_Bta14287 Protein
MHVYLLSAVAISIFLNFQEAVGWKATIYDDVQHGKTLQTLSGGPCQEVKDANNDRASSIN
TWGGCVVIYSDFQCKGTNKTMRPGSPHHKDFKLLHFDNVVSSIGPCP

>G2_Bta11629_BTA023203_2 cDNA
ATGGTCGCCTTCAGAGACGCAGTGCAGGTGGCTGCAGTTCTCGCTTTTATCTTGACACAA
TGCACTTCAGGGGAGGAGGCGACGGCGAGGCAGAAGCGTGGCGGTTTGGCGATGCTTAGC
TTCGGGCAGAAGCAGCAGGTCGTCCAGGAGACGGAAACGGTGACGACAACGACGACGACG
AAGGTGGTGCGGCGGAACGACGTCTTCGGGTACAACGAGCGGATGAGCAAGCCGCCGCCG
CCGGTCTCAGTGTGTCGCCGCTACGGCTACGACTCCGAGGGCCTCTACTCCATCAAGGCC
TTCACGAAGCTCCTGGAGAGCCTGAATAAGGCCCAGTGCGACATCGTCCAGGAGATCATC
TACCCGGCCGACAACAGCCTGAGCCTCCGGGACGACGACACCGCCGACTACATCGTCGTC
GGCCTCGGGGTCGCCGGCTCCGTCATCGCCAGTCGCCTCTCCGAGATCCCCTCCAACTCC
GTCATCGGTCTCGAGGCTGGTGGCAATCCCAGCTTGATCACCGAGTTCCCCAGGCTATGG
ACAGAGGCCGCAGACACGGAATACGAATGGAACTACCGCTTGAGGGCGGACAACTACAGC
TGCCTATCATCCGAGTACCAGCGCGGGAAGCTGTACAAAGGCCGCGGTCTGGGCGGCACT
TCCGCGATCGACCTGATGATCTACGACCGTTGCATGACATCCGACTACGAGAAATTCAAC
GAAATCGGTCTCCCGGACTGGGGCTACGAAGCCTGCCTGAAATACTACATGAAATCGGAG
GACGCCAGGTGCGAGGAGATCTTCACCAAGGTGACGACCATCCGGGCGTCGCACAACTCC
GGAGGCCTCCTCACCGTCGACTCCTTCTACAACAGCCAGACCGTCTCCATCCGCAAGTCG
TTCGGCCAGTGCATCTCCGAGCTCGGGATGACGAGCAAGGACGTCTTCACCGAGATCGAC
CACGACGGGTTCGCCCCCTCGTTGGCGCTCATCAAGGACGGTCTCCGTGTCAACGTGGCC
AGGGCCTTCCTGAGCCCGCGGGAGGTGCGGCGCCGACCCAACTTGAAAGTCTGCAAGTTC
TCCTACGTGACGAGGCTTCTCCTCGACGATGCCAACACCAAAGTCATCGGCGTCGAGTTC
CGGAATAGGCTCGGCGACCTGGTCAAGTTGTACTGCAACAAGGAGGTGATCCTGACGGCC
GGCGCTATCGAGTCCCCGCACGTCTTGCAGAACTCCGGCATCGGCGACCCGGTCGACTTG
GCGCGCGTCGGCACCATCCTCCGCAAAGAGCGTCCCGGCGTCGGGAAGAACCTCCAGATG
CACCCGCTCTTCCCGGGCGCCATCATGACGTTCAAGACGCCGCCGATCGCCCAGTGGCCG
GTCGGGGACATGCTCTACGAGCTCATCGCCCACCGCAAGGGCCCCATCACCAACATCGGA
CTCACCAGCTACACCGGCTTCGTCAAGACCAACACGAGCGACGTCCCGAACCTCCAGATC
ATCCCGTACTACTTCTCCAAGGAGGACTCTCTCTTCCTCAACGGGCAGCTCGCCGTCTTC
GGCTACGACGAGGCCACCCGCAAGAAGATCCGCGAGATCAACCGCGAGAGCTCCATCGCC
ATGTTCGGCGTGTCGCTCTTGTACCCCAAGTCCGTGGGCTCCGTCTACCCGGAAACGAGC
AACCCCTACGACCACCCGGTCATCGACGCTAAGCTCTTGAGCGACCCGGAGGACATCTAC
TCCGTCTTGGGAGGAATCGACTGGGTCCGAGGCCTGGAGGATTCCCCCGTGTTCCAGAAA
TTCGGCGCCAAGTTCCTACCGCTGTATGTCGACAACTGTCTCTACCCTATCGGCTCTAGG
GAGTACTGGATGGAGGCTGTCAGACACCTGACAGTTCCGCGGAAGCACCCTGTCGGCACG
TGTAAGATGGGTCTGCCCGACGATCCGCTGGCAGTCGTCGATCCTGATTTCCGGTTCATC

GGACTCGATAACCTCCGCATTGGTGACTCCTCCATCTTGCCCAGGATATTCTCGTCCGAC
TCCAGCGCTGTTACCGTCATGATCGCTGAGAAATGCGCTGATAGAATCATGAGTCAAATT
TGA

>G2_ Bta11629_ BTA023203_2 Protein
MVAFRDAVQVAAVLAFILTQCTSGEEATARQKRGGLAMLSFGQKQQVVQETETVTTTTTT
KVVRRNDVFGYNERMSKPPPPVSVCRRYGYDSEGLYSIKAFTKLLESLNKAQCDIVQEII
YPADNSLSLRDDDTADYIVVGLGVAGSVIASRLSEIPSNSVIGLEAGGNPSLITEFPRLW
TEAADTEYEWNYRLRADNYSCLSSEYQRGKLYKGRGLGGTSAIDLMIYDRCMTSDYEKFN
EIGLPDWGYEACLKYYMKSEDARCEEIFTKVTTIRASHNSGGLLTVDSFYNSQTVSIRKS
FGQCISELGMTSKDVFTEIDHDGFAPSLALIKDGLRVNVARAFLSPREVRRRPNLKVCKF
SYVTRLLLDDANTKVIGVEFRNRLGDLVKLYCNKEVILTAGAIESPHVLQNSGIGDPVDL
ARVGTILRKERPGVGKNLQMHPLFPGAIMTFKTPPIAQWPVGDMLYELIAHRKGPITNIG
LTSYTGFVKTNTSDVPNLQIIPYYFSKEDSLFLNGQLAVFGYDEATRKKIREINRESSIA
MFGVSLLYPKSVGSVYPETSNPYDHPVIDAKLLSDPEDIYSVLGGIDWVRGLEDSPVFQK
FGAKFLPLYVDNCLYPIGSREYWMEAVRHLTVPRKHPVGTCKMGLPDDPLAVVDPDFRFI
GLDNLRIGDSSILPRIFSSDSSAVTVMIAEKCADRIMSQI

>G3_bta08713_BTA007921.1 cDNA
ATGAAAGTGGTATTATTGTTGGGGCTTGTGGCCTTTGCCGCGGTGTACATCCAACCGAGC
GAGGCATGTGGAGAGTGTGACTCATGTTGGAATTACAAACACTGTGAAATAACCGAGTTT
CCCTGGGTTGTGAGGAGGTACTATGCCGTTTTCATAACCAGAGTAACCCTCAACAATTTC
TGGGGCAAAGGAAATTCTTGCGGAATGTTCAACAAATGTTCGGCTAAGGCTTGCGGCTGC
ATTGGATTAGACGTTGGCCGAATCGGAGGGCGGACATTGCAAGGACAATGGTTTTCCTAC
CACAGGAAGAAGCACATACAGTACATGCGTCGATTCTGGGCCGTTGTTAACGAAAACAAT
GCTGAGATGACTTTCTTCTTAGCCGCCGAGGCCTCGTGGAAGGTACCAGTGACAGTAGTA
AAAAAAGTAGCGTGTGACGAATGTGGAACGGTCCGCATAGAAACATACGTTGTACCGGAC
ACAGAGGTGAAAGTAAGAACTGTGTCCGTCGAGTTCAGAGCTATCATGCTTATCTTAAAG
ATTAAGTTTAGGATATTGTTGATGTGCAATGGCCTGAAAGGAGTTGTCAGTAACAAGCCG
TTCGCACTTGTTTTGGCTACGATGCCGTTCCCGGGTAAGAGCGACACGTGGGTCAAGATC
AACGCCGCCTTGCGCCATTACAACTTCGATCCGCACAAGATGATGTTCTTGAGGCACGAG
GCCTGCATCTACCCGCCCTTCGATTGGGCCAACTCCTGGGTCGGGCCTGCTAACAACAAC
TGCAACATTTGCAAAATTGACAAGTAG

>G3_bta08713_BTA007921.1 Protein
MKVVLLLGLVAFAAVYIQPSEACGECDSCWNYKHCEITEFPWVVRRYYAVFITRVTLNNF
WGKGNSCGMFNKCSAKACGCIGLDVGRIGGRTLQGQWFSYHRKKHIQYMRRFWAVVNENN
AEMTFFLAAEASWKVPVTVVKKVACDECGTVRIETYVVPDTEVKVRTVSVEFRAIMLILK
IKFRILLMCNGLKGVVSNKPFALVLATMPFPGKSDTWVKINAALRHYNFDPHKMMFLRHE
ACIYPPFDWANSWVGPANNNCNICKIDK

>G4_Bta08784_BTA021638.1 cDNA
ATGAGAACGTTTCTAGGAATACTTTGTGTGACCCTTGCAATCCAGCTGGTGATCGTCTTC
AGCGATGCTGAGCAGCATGTCCCTGGGCGATTCAAGGCTGGCCTAGACTTCCACCAAGAG
GCTGGTGACAAGGCCTCAACCCGCTCCATCACCTCCTCGGAACCCGGCGCCGAATGGGCT
AAGAGCTCCGTGCAAACCCCGATCGAAGATGAGATCACCGACTTGAACAAGGAGAAGGAA
GCCAGCAAGAAGAAGGAGGCCAATGCTGAGGCCGGCGAGGACCTCGAGGACCTTGAGGAT
TTGGAGGATATCGATCACCTGGCCACCCGTGGTCTCGAAGACTTGGACGAGGATTTGGGG
GAGGATGATTTGGATGTGGAAGGAGCCGCCGACCATGAACTTGAAACACGCGGACTCCTT
GAATCAGGAATGAACATGCTCCCGTCGCGCGAGGCCTTGAGTCATACCGTCTCCGACCTG
AAGGACAGGGCCGCGAGTATCCCGAGGGCCGTTTCGGACATGTACCACGACCCGCAGGCC
GCCCGTGAATCCCTCCGCAACACCCTGAGCAACGCCAAGGACACCGTCATGAACTCAAAC
GTTCGCGCCAAGATGGCCGATCTCTACAAGGCCGGACTCGACAAGGGCCTCGACCTCTAC
AAGGGCATCAACCAGCGTCTCAGCCCCGGCATGGCCAAGACCCTCAACCAAGGCGTCAGC
TCCACCATGCGCACCATCCAAGCCGCCAAGGAGTCCCCCATCGCCAAGTCCATCTCCGAG
AAGGTCGCCGCCTGGTTGCCCTGA

>G4_Bta08784_BTA021638.1 Protein
MRTFLGILCVTLAIQLVIVFSDAEQHVPGRFKAGLDFHQEAGDKASTRSITSSEPGAEWA
KSSVQTPIEDEITDLNKEKEASKKKEANAEAGEDLEDLEDLEDIDHLATRGLEDLDEDLG
EDDLDVEGAADHELETRGLLESGMNMLPSREALSHTVSDLKDRAASIPRAVSDMYHDPQA

ARESLRNTLSNAKDTVMNSNVRAKMADLYKAGLDKGLDLYKGINQRLSPGMAKTLNQGVS
STMRTIQAAKESPIAKSISEKVAAWLP

>Solyc05g015390.2.1 REF-like stress related protein 1 cDNA
ATGTCTACCAACAAGGTTGAGATGGAGAAATGCGAGGTTAATTTGAAACATCTAGGGTTT
GTAAGGGTGTTGGCTCTAAACACTGCAGTTTTGTTGTCAAATTTGTATGATTATGCGAAG
CAGAACTCGGGGCCTCTGAGATCAACTGTTGGTACAGTAGAAAACGCTGTAACCACCGTC
GTAAGGCCTGTTTACGATAGATTTAAAGGTGTTCCTGATGAAGTCCTTGTTTTCCTAGAC
AAAAAGTTGGATGATGGAACAGCAAAATTTGATGAACATGCTCCTCCATTGGCCAAGAAG
GTTGTCAGCAGTGTCCAGTCTTTGTTTCAGAAGGCATCAGAGGTAGCACAAGGCTTGTCC
AAAGACATACAGGAGGGAGGTCCTCGTGCAGCTATCTATCATGCTGGTGAATTGTCTAAG
CAATTTGGTACTACGCAAGTGGCAGTACTCTGGTATCATGTCAATCATTGTCCACCATTG
CACGGAATCGCACAGATGGCTGCTCCTACAGTTGCTCACTGGTCCGAAAAGTACAATCAT
TTGGTAGCTAACTTGCAAGAGAAGGGTTATAGTGTTGTCAGCTATATCCCTTTGATACCT
GTTGAAGAAATCTCAAAGGCATATAAACAGGTTGAATCTGCTGCAACGAAGAAAGAAGGT
GCCACTGACTCGACCTCTAGTAAATCTGAATGA

>Solyc05g015390.2.1 REF-like stress related protein 1 Protein
MSTNKVEMEKCEVNLKHLGFVRVLALNTAVLLSNLYDYAKQNSGPLRSTVGTVENAVTTV
VRPVYDRFKGVPDEVLVFLDK**KLDDGTAKFDEHA**PPLAKKVVSSVQSLFQKASEVAQGLS
KDIQEGGPRAAIYHAGELSKQFGTTQVAVLWYHVNHCPPLHGIAQMAAPTVAHWSEKYNH
LVANLQEKGYSVVSYIPLIPVEEISKAYKQVESAATKKEGATDSTSSKSE*

>Solyc01g099770.2 meloidogyne-induced giant cell protein DB141 cDNA
ATGTTGGTTTATCAGGATCTCCTCACCGGTGATGAGCTTCTCTCTGACTCATTTCCCTAC
AAAGAAGTCGAGAATGGCGTGCTTTGGGAAGTTCAAGGGAAGTGGGTTGTTCAAGGTGCC
GTCGATGTAAACATTGGGGCAAATCCTTCTGCTGAGGGTGGATGTGAAGATGAAGGTGTG
GATGACCAAGCTGTCAGGGTTGTCGATATTGTTGACACTTTTAGACTTCAGGAGCAACCT
GCTTTTGACAAGAAACAATTTGTTACCTTCATGAAGAGATACATCAAGAACCTGACACCC
AAGCTAGAAGGAGAAACACAAGAAGCATTTAAAAAGAACATTGAAGCAGCAACTAAGTTT
CTTTTGCAAAAGATCAAGGACTTGCAATTCTTTGTTGGTGAGAGCATGCATGATGATGGC
GCCCTGGTGTTTGCGTACTACAAGGAGGGTTCAGCTGATCCTACCTTTTTGTACATTGCA
CCTGGTTTGAAGGAGATCAAGTGCTAG

>Solyc01g099770.2 meloidogyne-induced giant cell protein DB141 Protein
MLVYQDLLTGDELLSDSFPYKEVENGVLWEVQGKWVVQGAVDVNIGANPSAEGGCE**DEGV**
**DDQAVRVVDIVDTFRLQEQPAFDKKQFVTFMKRYIKNLTPKLEGETQEAFKKNIEAATKF**
**LLQKIKDLQFFVGE**SMHDDGALVFAYYKEGSADPTFLYIAPGLKEIKC*

>Solyc01g099780.2.1 Translationally-controlled tumor protein homolog
ATGTTGGTTTATCAAGATCTCCTCACTGGTGATGAGCTTCTCTCAGACTCATTTCCCTAC
AAAGAACTCGAGAATGGAATGCTTTGGGAAGTTCAAGGGAAGTGGGTTGTTCAAGGTGCC
GTTGATGTAAACATTGGGGCAAATCCTTCTGCTGAGGGTGGAGGTGAAGATGAAGGTGTG
GATGACCAAGCTGTCAAGGTTGTCGATATTGTTGACACTTTTAGACTTCAGGAACAACCT
GCTTTTGACAAGAAACAATTTGTTACCTACATGAAGAGATACATCAAGAACCTGACACCC
AAGCTAGAAGGAGAAACACAAGAAGCATTTAAAAAGAACATTGAATCAGCAACTAAGTTT
CTTTTGCAAAAGATCAAGGACTTGCAATTCTTTGTTGGTGAGAGCATGCATGACGACAGT
GCCCTGGTGTTTGCGTACTACAAGGATGGTTCAGCTGATCCTACCTTTTTGTACCTTGCA
CCTGGCTTGAAGGAGATCAAGTGCTAG

>Solyc01g099780.2.1 Translationally-controlled tumor protein homolog 1
MLVYQDLLTGDELLSDSFPYKELENGMLWEVQGKWVVQGAVDVNIGANPSAEGGGEDEGV
DDQAVKVVDIVDTFRLQEQPAFDKKQFVTYMKRYIKNLTPKLEGETQEAFKKNIESATKF
LLQKIKDLQFFVGESMHDDSALVFAYYKDGSADPTFLYLAPGLKEIKC*

>Red Firefly luciferase Arabidopsis codon optimized 1654 bp
ATGGAGGACGCAAAAAACATAAAAAAAGGCCCAGCCCCCTTTTATCCCTTAGAAGACGGC
ACAGCCGGCGAACAACTGCACAAAGCAATGAAGCGATATGCTTTAGTTCCGGGAACCATC
GCATTCACGGATGCCCATATAGAGGTAAACATCACCTACGCGGAATACTTTGAAATGAGT
GTTCGATTGGCAGAGGCCATGAAACGATATGGACTCAATACCAATCACCGAATCGTAGTT
TGTTCTGAAAACAGCCTCCAATTTTTCATGCCTGTGCTTGGAGCATTATTTATCGGTGTT

GCCGTTGCCCCGGCCAATGATATATACAATGAGCGTGAACTGCTCAACAGTATGAATATC
TCTCAACCCACTGTTGTATTTGTGTCAAAAAAAGGGCTTCAGAAAATTCTCAACGTACAG
AAGAAATTACCAATAATCCAGAAGATAATCATAATGGACTCTAAAACTGATTACCAGGGC
TTCCAGTCCATGTACACATTTGTCACTTCTCATCTGCCCCCGGGATTCAACGAGTATGAC
TTCGTTCCGGAGAGCTTCGACAGAGACAAGACTATAGCTCTGATTATGAACAGCAGTGGA
TCTACTGGATTACCCAAAGGCGTGGCGCTCCCACACAGAGCTCTTTGTGTGCGATTCAGC
CACGCACGAGACCCTATATTCGGCAATCAGATAGCGCCTGATACCGCCATTCTTAGCGTA
GTTCCTTTTCATCACGGTTTCGGAATGTTCACGACGTTGGGTTATTTAATTTGTGGTTTT
CGAGTAGTTCTGATGTACAGGTTCGAAGAAGAATTGTTCCTACGATCCTTGCAGGACTAC
AAAATACAAACAGCTCTATTAGTCCCCACGCTCTTCTCCTTCCTGGCTAAATCAACCCCTC
ATAGACAAATATGATTTAAGCAACTTACACGAAATCGCAAGTGGTGGAGCGCCCCTGAGC
AAGGAGGTAGGGGAGGCAGTCGCAAAACGTTTCCACCTGCCGGGCATTAGGCAGGGCTAT
GGCCTTACTGAAACAACGTCCGCTATCCTGATTACTCCAAAAGGTGATGACAAACCAGGA
GCGGTCGGCAAAGTTGTCCCGTTCTTCGAAGCCAAGGTTGTGGACCTTGACACGGGAAAG
ACGCTCGGCGTAAATCAGCGTGGTGAATTATGTGTCAGAGGCCCGATGATTATGAGCGGC
TATGTAAATAATCCAGAGGCTACGAACGCACTCATTGACAAGGATGGTTGGCTGCACTCC
GGCGATTTAGCTTATTGGGATGAAGACGAGCACTTCTTCATTGTAGGCAGACTGAAGTCA
CTCATAAAATACAAGGGCTACCAGGTGGCTCCGGCAGAACTCGAGAGCATCCTTCTACAA
CACCCCAACATTTTCGACGCAGGGGTGGCAGGTTTACCAGATGATGATGCGGGGGAGCTC
CCTGCAGCGGTAGTAGTGCTCGAACATGGAAAAACGATGACGGAGAAAGAGATCGTGGAC
TACGTTGCCTCTCAGGTTACGACTGCTAAAAAGCTAAGAGGTGGTGTGGTTTTTGTTGAT
GAAGTACCAAAGGGCCTCACAGGTAAACGTGACGCAAGAAAGATACGAGAGATATTGATA
AAAGCTAAGAAGGGAGGCAAAAGCAAACTCTAG*

**Figure S1: Sequences of cDNA/ proteins used in this study.** Underlined: predicted signal peptide. grey: cDNA part in VIGS construct. Bold: overlapping regions between different prey in the Y2H assay (named i-part).

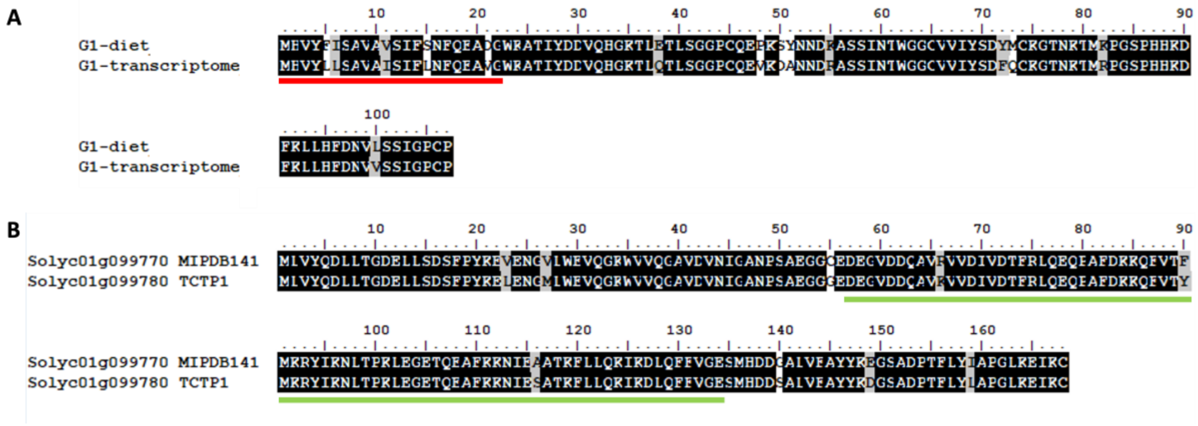


**Figure S2: Protein alignment G1 and TCTPs.** A: Alignment between G1 identified in the artificial diet and G1 identified in the transcriptome. Red underlined: predicted signal peptide. B: Alignment between MIPDB141 and TCTP1 from *Solanum lycopersicum*. Green underlined: overlapping regions between different prey in the Y2H assay (named i-part).

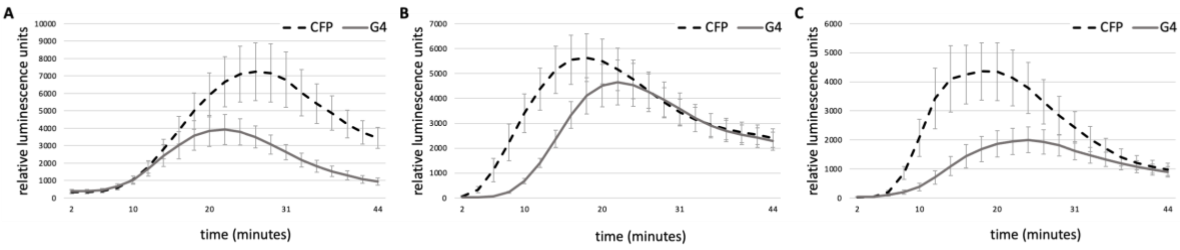

**Figure S3: ROS assays with effector G4 in different plant species to show production of ROS in time.** A: ROS production in *Nicotiana benthamiana.* B: ROS production in *Solanum lycopersicum*. C: ROS production in *Capsicum annuum*. Data and statistical analysis presented in Figure 2.

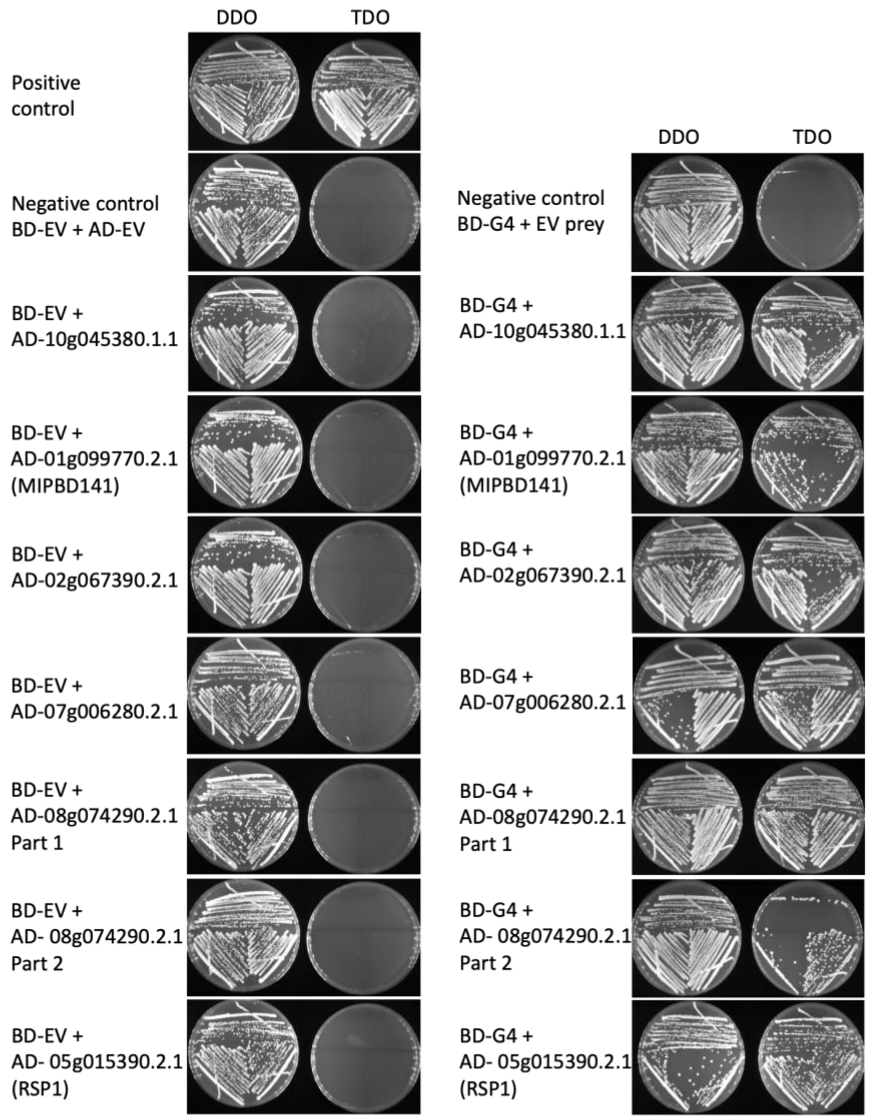


**Figure S4: confirmation of interaction G4 with truncated protein of a selection of candidate target proteins using yeast two hybrid 1-by-1 assays.** Co-transformation of the selected targets of effector G4 Selection was performed on DDO to test for transformation of both prey and bait constructs. TDO selection was performed to confirm interaction with G4 and exclude auto-activation.

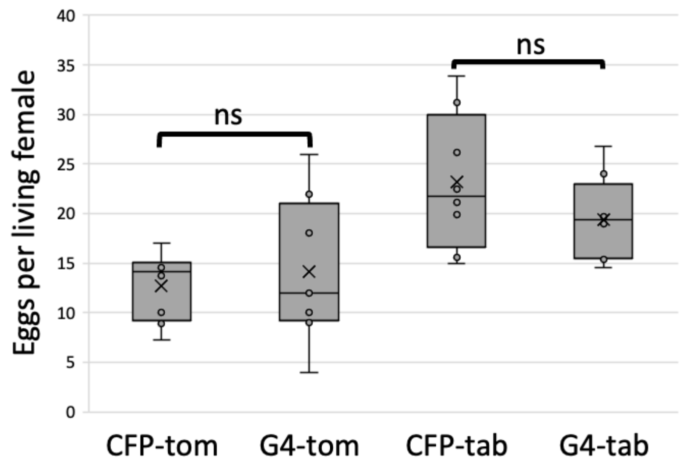


**Figure S5: Whitefly bioassay on *Solanum lycopersicum* (Moneymaker) and *Nicotiana tabacum* transiently expressing G4.** CFP: free CFP as control. ns indicates no significant difference compared to the CFP control shown by paired T-test. Graphs represents the eggs per living female of 8 plants.

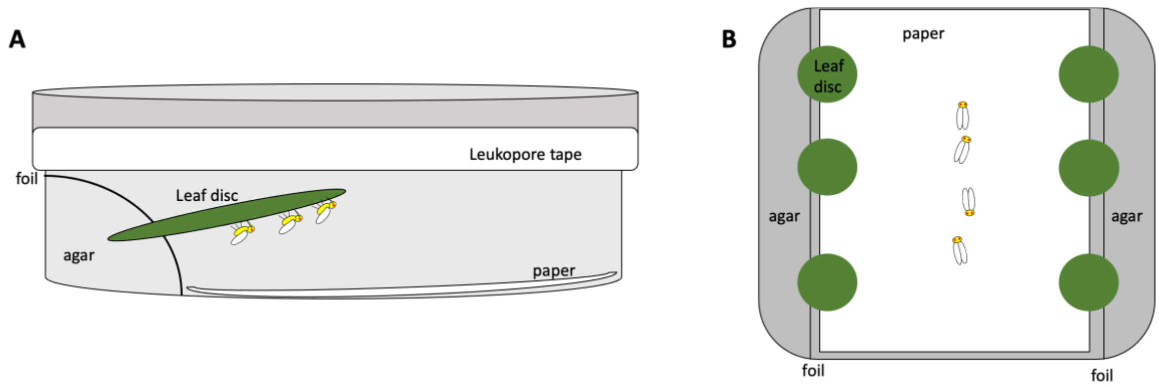


**Figure S6: Setup VIGS whitefly bioassays.** A: No-choice bioassays with either one GFP-silenced or one MIPBD141/TCTP-silenced leaf discs. B: Choice assay with on one side three GFP-silenced leaf discs and on the other side three MIPBD141/TCTP-silenced leaf discs.

**Table S4:** Primer list

| Name | 5' --> 3' sequence |
|---|---|
| REDFF_NLUC_GWY_FW | CTAGTCTAGAACCATGATAGATTACAAAGACGACGATGACAAGGAGGACGCAAAAAACATAA |
| REDFF_NLUC_GWY_RV | CTAGTCTAGACCTCCACCGGATCCGCCCATAATCATCGGGCCTC |
| REDFF_CLUC_GWY_FW | CTAGTCTAGAACCATGATATACCCTTACGATGTGCCTGACTACGCTGGCCCGATGATTATGAG |
| REDFF_CLUC_GWY_RV | CTAGTCTAGACCTCCACCGGATCCGCCGAGTTTGCTTTTGCCTC |
| GWY_REDFF_NLUC_FW | CTAGAGCGCTGGCGGATCCGGTGGAGAGGACGCAAAAAACATAA |
| GWY_REDFF_NLUC_RV | CTAGAGCGCTCTACTTGTCATCGTCGTCTTTGTAATCCATAATCATCGGGCCTC |
| GWY_REDFF_CLUC_FW | CTAGAGCGCTGGCGGATCCGGTGGAGGCCCGATGATTATGAG |
| GWY_REDFF_CLUC_RV | CTAGAGCGCTCTAAGCGTAGTCAGGCACATCGTAAGGGTAGAGTTTGCTTTTGCCTC |
| G1-UTR-F | CCTGAGTATTCTGCAAACTC |
| G1-UTR-R | CAATAGCCTGCGATATATCC |
| G1attb1No-SPstart | aaaaagcaggcttaATGTGGAAGGCGACAATTTATGATG |
| G1attb2stop | agaaagctgggtgTTATGGGCAGGGACCGATG |
| G3_XP_018898154_F_FL | ATGAAAGTGGTGGTTGCTGTTG |
| G3_XP_018898154_R_FL | CTACTTGTCAATTTTGCAAATG |
| G3attb2stop | agaaagctgggtgCTACTTGTCAATTTTGCAAATG |
| G3attb1start | aaaaagcaggcttaATGAAAGTGGTATTATTGTTGG |
| G4_Bta08784_R_FL | TCAGGGCAACCAGGCGG |
| G4_Bta08784_F_FL | ATGAGAACGTTTCTAGGAATAC |
| G4_attb1_F_SP+start | aaaaagcaggcttaATGAGAACGTTTCTAGGAATAC |
| G4_attb1_F_noSP+start | aaaaagcaggcttaATGGAGCAGCATGTCCCTGG |
| G4_attb2_R_stop | agaaagctgggtgTCAGGGCAACCAGGCGG |
| G4_attb2_R_nostop | agaaagctgggtgGGGCAACCAGGCGG |
| attb short 1 | ACAAGTTTGTACAAAAAAGCA |
| attb short 2 | ACCACTTTGTACAAGAAAGCT |

**Table S5:** List of modified destination vectors used in the luciferase complementary assay.

| ID | Gene name | Backbone | Template/source | Procedure |
|---|---|---|---|---|
| bgIFP 4425 | P19 inhibitor silencing | pBIN61 | | |
| bgIFP 5593 | HA-RedFFLuc Cterm-Gateway box | pGWB402 | | |
| bgIFP 5594 | Gateway box - RedFFLuc Nterm-FLAG-Stop | pGWB402 | | |
| bgIFP8186 | Rep TY Alb13 - RedFF Nterm FLAG GWB402 | pGWB402 | bgIFP5594 | Gateway |
